# TORC1 regulates the transcriptional response to glucose and developmental cycle via the Tap42-Sit4-Rrd1/2 pathway in *Saccharomyces cerevisiae*

**DOI:** 10.1101/859793

**Authors:** Mohammad Alfatah, Jin Huei Wong, Vidhya Gomathi Krishnan, Yong Cheow Lee, Quan Feng Sin, Corinna Jie Hui Goh, Kiat Whye Kong, Shawn Hoon, Prakash Arumugam

## Abstract

Target of Rapamycin Complex 1 (TORC1) is a highly conserved eukaryotic protein complex that couples the presence of growth factors and nutrients in the environment with cellular proliferation. TORC1 is primarily implicated in linking amino acid levels with cellular growth in yeast and mammals. Although glucose deprivation has been shown to cause TORC1 inactivation in yeast, the precise role of TORC1 in glucose signaling and the underlying mechanisms remain unclear. In this paper, we demonstrate that the presence of glucose in the growth medium is both necessary and sufficient for TORC1 activation. TORC1 activity increases upon addition of glucose to yeast cells growing in a non-fermentable carbon source. Conversely, shifting yeast cells from glucose to a non-fermentable carbon source reduces TORC1 activity. Analysis of transcriptomic data revealed that glucose and TORC1 co-regulate about 27% (1668/6004) of yeast genes. We demonstrate that TORC1 orchestrates the expression of glucose-response genes mainly *via* the Tap42-Sit4-Rrd1/Rrd2 pathway. To confirm TORC1’s role in glucose-signaling, we tested its role in spore germination, a glucose-dependent developmental state transition in yeast. TORC1 regulates the glucose-responsive genes during spore germination and inhibition of TORC1 blocks spore germination. We propose that a regulatory loop that involves activation of TORC1 by glucose and regulation of glucose-responsive genes by TORC1, mediates nutritional control of growth and development in yeast.

## INTRODUCTION

Cells sense changes in nutrient availability in their environment and accordingly adjust their growth and developmental cycles. Glucose-response and spore germination in *Saccharomyces cerevisiae* are excellent model systems to study this biological phenomenon. When glucose is added to yeast cells growing in a non-fermentable carbon source, the transcriptome and metabolome are extensively reprogrammed to facilitate their growth in the new milieu. This adaptation is referred to as the ‘transcriptional response to glucose’ (will be referred to as glucose-response here). Likewise, when haploid spores are transferred to a nutrient medium that contains glucose (or a rapidly fermentable carbon source), they exit their state of ‘hibernation’ and re-enter the mitotic cell cycle (*1*). Exactly how changes in glucose levels in the environment are sensed by the cell leading to dramatic modulation of growth and developmental regulatory circuits is poorly understood.

In yeast, glucose levels in the environment are thought to be mainly sensed by the cyclic AMP (cAMP)-dependent protein kinase A (PKA) (*2*). Addition of glucose or a rapidly fermentable sugar to the medium activates PKA *via* GTP-binding proteins Ras1/2 and Gpa2 which results in activation of adenylate cyclase and cAMP production (Figure 1). PKA consists of a catalytic subunit (Tpk1, 2 or 3) and a regulatory subunit Bcy1. Bcy1 inhibits PKA’s catalytic activity and this inhibition is relieved by binding of cAMP to Bcy1 causing its dissociation from the catalytic subunit (Figure 1). PKA phosphorylates several cellular proteins leading to enhanced protein synthesis and inhibition of stress response.

**Figure 1.**
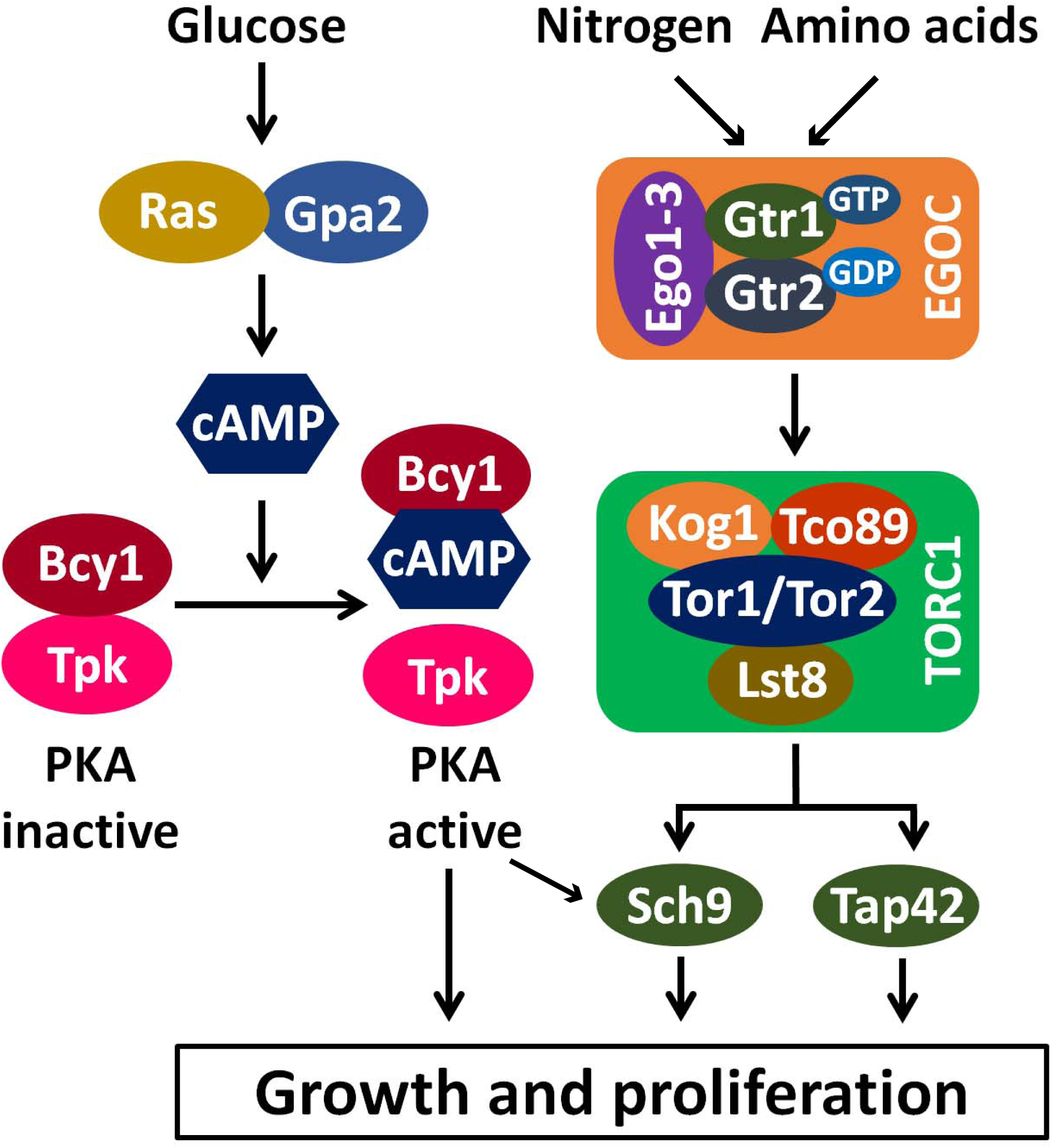
PKA and TORC1 connect the presence of glucose and amino acids/nitrogen levels respectively with cell proliferation in yeast. See the introduction for details.

While glucose signaling is mainly attributed to PKA, the coupling of amino acid levels and quality of nitrogen source with cellular growth is performed by TORC1, a multi-subunit protein complex conserved among eukaryotes (*2, 7*). TORC1 promotes anabolic processes such as protein synthesis and inhibits catabolic processes like autophagy. TORC1 in *Saccharomyces cerevisiae* is composed of four subunits, namely Tor1/Tor2 (a serine-threonine kinase), Lst8 (equivalent of mLst8/GβL), Kog1 (equivalent of mammalian raptor) and Tco89 (a yeast-specific subunit). The conserved Rag GTPases Gtr1 and Gtr2, act upstream of TORC1 and link the amino acid and nitrogen levels with TORC1 activity (Figure 1). Gtr1 and Gtr2 form a heterodimer and the state of the nucleotides bound to Gtr1-Gtr2 determines whether they activate or inhibit TORC1. Specifically, a heterodimer of Gtr1/Gtr2 in which the Gtr1 is bound to GTP and Gtr2 is bound to GDP, activates TORC1. Binding of Gtr1/Gtr2 to TORC1 is facilitated by three proteins Ego1/Meh1, Ego2 and Ego3/Slm4, which together with Gtr1/Gtr2 constitute the EGO complex. TORC1 links changes in nutrient levels with transcriptomic reprogramming *via* its downstream effectors, namely Sch9 (a serine-threonine kinase) and Tap42 (a PP2A phosphatase-binding protein).

During the glucose response, yeast cells reshape their metabolism by switching from oxidative phosphorylation to glycolysis to obtain energy. About 40% of yeast genes are altered in their expression upon glucose addition and 90% of these transcriptional changes could be induced by activation of either Ras2 or Gpa2 (*14*). Furthermore, an activated form of PKA recapitulated 90% of transcriptomic changes induced by glucose indicating that PKA is the main regulator of the transcriptional response to glucose in yeast (*15*).

The role of TORC1 in regulating the glucose-response is not clear. A transcriptomic study showed that PKA works along with TORC1 in regulating the transcriptional response to glucose (*16*). However, another study found that the major TORC1 effector Sch9 plays a very minor role in the glucose response (*15*). Although, activation of TORC1-effector kinase Sch9 recapitulated a number of transcriptional changes induced by glucose, inactivating Sch9 had only a minor effect on the glucose-response (*15*). It is also unclear whether the role of Sch9 in glucose-response is downstream of TORC1 activation or independent of TORC1 activity. Thus, it is very important the clarify TORC1’s precise function in the glucose-response.

In this paper, we show that TORC1 activity is upregulated during the glucose-response. TORC1 is activated by glucose through Gtr1/Gtr2-dependent and Gtr1/Gtr2-independent mechanisms. Transcriptomic analysis revealed that about 50% of glucose-response genes are regulated by TORC1. We show that TORC1 activity is required for establishment and maintenance of transcriptional response to glucose. TORC1 regulates the glucose-response genes through its downstream effectors Tap42/ Sit4/Rrd1-Rrd2. Our results are consistent with the model that inhibition of TORC1 leads to activation of Sit4/Rrd1-Rrd2 phosphatase which effects changes in the expression of glucose-response genes. If TORC1’s role in glucose-response is physiologically relevant, then TORC1 inhibition affect spore germination, a glucose-dependent developmental state transition in yeast (*1*). This prediction was confirmed by our observations that TORC1 regulates the glucose-responsive genes during spore germination and is essential for spore germination. We propose that TORC1 is an important regulator of the glucose-response and this function is essential for developmental state transition from quiescent spores into actively growing vegetative cells.

## RESULTS

### Glucose is necessary and sufficient for TORC1 activation

To investigate the role of TORC1 in glucose signaling, we first tested whether glucose is required for TORC1 activation. To assess the kinetics of TORC1 activation, we tagged the TORC1 substrate Sch9 with 6 copies of the hemagglutinin (HA) epitope. Phosphorylation of Sch9 can be assayed by cleaving it with NTCB (2-nitro-5-thiocyanatobenzoic acid) followed by detecting the electrophoretic mobility of the C-terminal HA-tagged Sch9 fragment by Western analysis (*17*). We transferred log-phase wild type and *gtr1*Δ cells grown in synthetic medium containing 2% glucose (SC/D) into SC medium lacking glucose (SC-D). After 60’ following transfer, Sch9 was completely dephosphorylated in both wild type and *gtr1*Δ cells indicating that TORC1 activity requires the presence of glucose in the medium (Figure 2A). We transferred the glucose-starved cells back into SC medium-containing glucose (SC/D). TORC1 was reactivated immediately upon addition of glucose in the wild type strain (Figure 2A). However, TORC1 activation in the *gtr1*Δ strain was delayed by about 10 minutes in comparison to the wild type strain (Figure 2A). These results suggest that glucose activates TORC1 via Gtr1/2-dependent and Gtr1/2-independent pathways in yeast,

**Figure 2.**
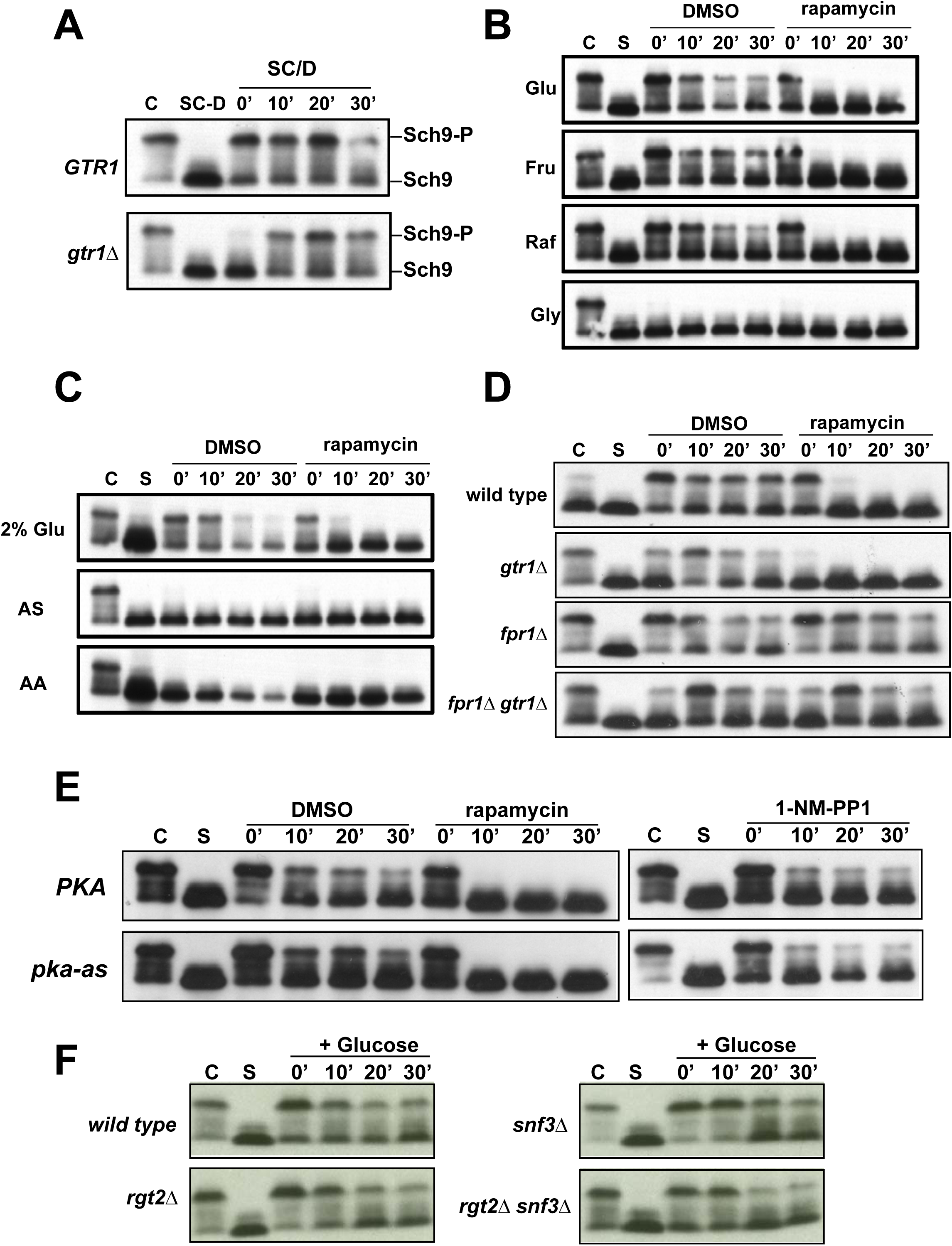
Presence of glucose in the medium is necessary and sufficient for TORC1 activation. A) Log-phase wild type and *gtr1Δ* cells grown in Synthetic medium with 2% glucose (SC/D) were transferred into Synthetic medium medium lacking glucose (SC-D) and incubated for 1 hour. Glucose-starved cells (SC-D) were then transferred back into SC medium (SC/D). Aliquots of the yeast cultures were taken after 0’, 10’, 20’ and 30’ and used for preparing protein extracts. Phosphorylation of Sch9 was monitored by Western blotting. B) Wild type cells in logarithmic phase (C) were subjected to complete nutrient starvation by incubating them in 0.3 M sorbitol for 1 hour. Starved cells (S) were then transferred to a solution containing either 110 mM glucose or 110 mM fructose, or 110 mM raffinose or 110 mM glycerol in the presence and absence of rapamycin (2 µM). Aliquots of the cultures were taken after 0’, 10’, 20’ and 30’ and used for preparing protein extracts. Phosphorylation of Sch9 was monitored by Western blotting. C) Wild type cells in logarithmic phase (C) were subjected to complete nutrient starvation by incubating them in 0.3 M sorbitol for 1 hour. Starved cells (S) were then transferred to a solution containing either 110 mM glucose or ammonium sulphate or amino acid mixture in the presence and absence of rapamycin (2 µM). Aliquots of the cultures were taken after 0’, 10’, 20’ and 30’ and used for preparing protein extracts. Phosphorylation of Sch9 was monitored by Western blotting. D) Wild type, *gtr1Δ*, *fpr1Δ* and *fpr1Δ gtr1Δ* cells in log phase were subjected to complete nutrient starvation by incubating them in 0.3 M sorbitol for 1 hour. They were then transferred to a 2% glucose solution in the presence and absence of rapamycin (2 µM). Aliquots of the cultures were taken after 0’, 10’, 20’ and 30’ and used for preparing protein extracts. Phosphorylation of Sch9 was monitored by Western blotting. E) Wild type and *pka-as* cells subjected to complete nutrient starvation were transferred to 2% glucose solution in the presence of either DMSO or rapamycin (2 µM) or 1-NM-PP1 (25 µM). Aliquots of the cultures were taken after 0’, 10’, 20’ and 30’ and used for preparing protein extracts. Phosphorylation of Sch9 was monitored by Western blotting. F) Wild type, *rgt2Δ*, *snf3 Δ*, and *rgt2Δ snf3Δ* cells subjected to complete starvation were transferred to 2% glucose solution. Aliquots of the cultures were taken after 0’, 10’, 20’ and 30’ and used for preparing protein extracts. Phosphorylation of Sch9 was monitored by Western blotting.

We then tested whether glucose is sufficient for TORC1 activation. We subjected log-phase yeast cells to complete nutrient starvation by washing off all the nutrients and transferring them into 0.3 M sorbitol. TORC1 was inactive after 1 hour of incubation in 0.3 M sorbitol (Figure 2B). We then added either glucose (110 mM) or equimolar amounts of fructose or raffinose or glycerol to starved cells, in the presence and absence of TORC1 inhibitor rapamycin (2 µM).

Addition of glucose or fructose or raffinose caused an immediate phosphorylation of Sch9 which was abolished by addition of rapamycin (2 µM) (Figure 2B). However, addition of glycerol did not activate TORC1. Unlike glucose, both ammonium sulphate and a mixture of all amino acids failed to activate TORC1 in the assay (Figure 2C). Our results indicate that glucose (or a rapidly fermenting carbon source like fructose/raffinose) is sufficient for TORC1 activation.

We then tested whether glucose-induced TORC1 activation requires its upstream regulator Gtr1. Addition of glucose caused immediate phosphorylation of Sch9 in wild type cells but with a 10-minute delay in *gtr1Δ* cells (Figure 2D). Sch9 phosphorylation in both wild type and *gtr1Δ* cells was abolished by rapamycin (2 µM) (Figure 2D). Rapamycin binds to the peptidyl-prolyl cis-trans isomerase Fpr1 and Fpr1-Rapamycin complex inhibits TORC1 by binding to Tor1 kinase (*18*). To determine whether rapamycin at 2 µM specifically inhibits TORC1, we examined the effect of rapamycin on glucose-induced Sch9 phosphorylation in wild type and *gtr1Δ* strains lacking *FPR1* gene. As observed previously, there was a 10-minute delay in the onset of Sch9 phosphorylation in the *fpr1Δ gtr1Δ* strain in comparison to the *fpr1Δ* strain. However, addition of rapamycin had no effect on Sch9 phosphorylation in both strains confirming that TORC1 is the Sch9-phosphorylating kinase. In summary, our data indicate that glucose is sufficient to activate TORC1 through Gtr1-dependent and Gtr1-independent mechanisms in yeast.

### Glucose-induced TORC1 activation is independent of PKA and glucose sensors Snf2 and Rgt2

As PKA is the main regulator of the transcriptional response to glucose in yeast (*2*), we tested whether glucose-induced TORC1 activation requires PKA activity. The catalytic unit of PKA is encoded by three genes *TPK1-3* in yeast. We constructed an analogue-sensitive allele of PKA (*pka-as*) by deleting *TPK3* and introducing gatekeeper mutations namely *tpk1-*M164G and *tpk2-*M147G (*15*). 1-NM-PP1 inhibited the growth of the *pka-as* strain at 3.125 µM but not the wild type strain suggesting that it specifically inhibits PKA activity in the *pka-as* strain (Supplementary Figure S1A). To ensure complete inhibition of PKA in our experiments we used 1-NM-PP1 at 25 µM. Addition of 1-NM-PP1 to *pka-as* but not *PKA* cells, reduced PKA activity as detected by Western analysis using an anti-PKA substrate antibody (Figure S1B).

However, glucose-induced Sch9 phosphorylation in 1-NM-PP1-treated *PKA* and *pka-as* cells were comparable (Figure 2E). These results indicate that glucose-induced TORC1 activation is independent of PKA activity. We also tested whether glucose-induced TORC1 activation is dependent on glucose sensors namely Rgt2 and Snf3 that regulate the transport of glucose into yeast cells (*2*). Glucose-induced TORC1 activation was comparable in wild type, *rgt2Δ*, *snf3Δ* and *rgt2Δ snf3Δ* strains (Figure 2F) precluding a role for these glucose sensors in glucose-induced TORC1 activation.

### TORC1 is activated during glucose-response

When glucose is added to yeast cells growing in a medium containing a non-fermentable carbon source, several genes are induced or repressed which restructure the transcriptional and metabolic state of yeast (*14*). This phenomenon is referred to as the glucose response (21) and the differentially expressed genes constitute the glucose-responsive genes. We investigated whether TORC1 activity is altered during the glucose-response. We added glucose (final concentration of 2%) to wild type and *gtr1Δ* cells growing in a medium containing non-fermentable carbon sources ethanol and glycerol (SC/EG) and examined the consequences on TORC1 activity by assaying Sch9 phosphorylation. TORC1 activity was low in cells growing in the non-fermentable carbon source and increased rapidly upon addition of glucose (Figure 3A). The increase in TORC1 activity upon glucose addition was delayed in the *gtr1Δ* strain as observed previously in our glucose/total starvation assays (Figure 2A and 2D). We also performed the converse experiment in which we switched the carbon source of yeast cells from glucose to ethanol-glycerol. TORC1 activity was drastically reduced upon transfer from SC/D to SC/EG in both wild type and *gtr1*Δ strains (Figure 3B).

**Figure 3.**
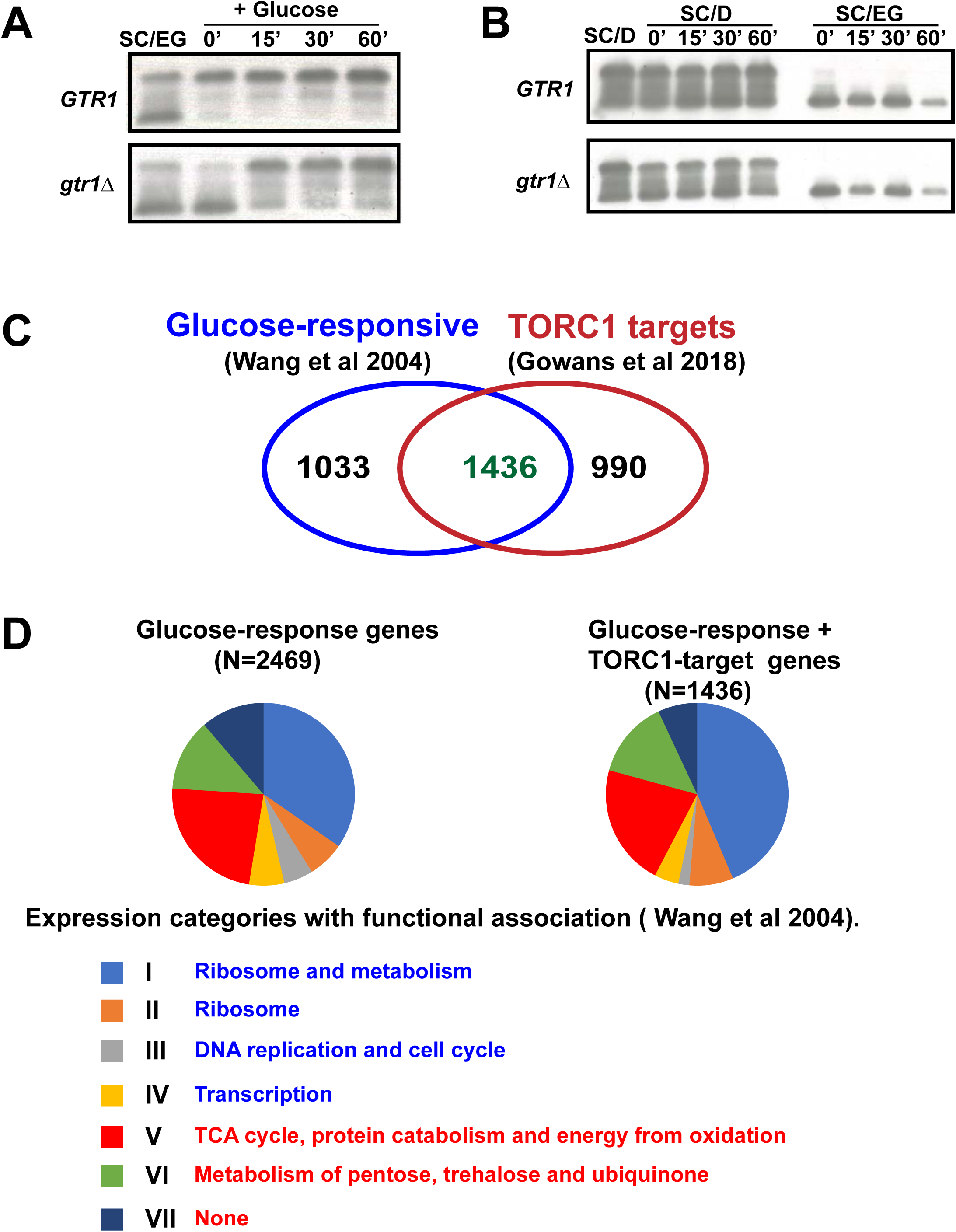
TORC1 is activated during glucose-response. A) Wild type and *gtr1Δ* cells were grown to logarithmic phase in SC/EG medium and then glucose (2% final concentration) was added to the cultures. Aliquots of the cultures were taken after 0’, 15’ 30’ and 60’ and used for preparing protein extracts. Phosphorylation of Sch9 was monitored by Western blotting. B) Wild type and *gtr1Δ* cells were grown to logarithmic phase in SC/D medium. Cultures were then divided into two parts. For one part, cells were pelleted and washed thrice with SC/EG medium, resuspended in SC/EG medium and incubated at 30 °C. The second part was transferred back to SC/D and was also incubated at 30 °C. Aliquots of the cultures were taken after 0’, 15’ 30’ and 60’ and used for preparing protein extracts. Phosphorylation of Sch9 was monitored by Western blotting. C) Venn-diagram showing the overlap of glucose-responsive genes (*14*) with TORC1 target genes (*19*). D) Pie-chart shown the distribution of the glucose-responsive genes and TORC1-glucose co-regulated (TGC) genes among the various gene clusters defined by response to glucose and Ras activation (Wang et al 2003). Functional enrichment among the various clusters induced and repressed by glucose are indicated, in blue and red fonts respectively.

### Overlap of TORC1 targets with the glucose-response genes

We then explored whether there is any overlap of glucose-response genes with TORC1 target genes reported in the literature. Based on the response to glucose and Ras activation, glucose-responsive genes (N= 3273) were classified into 8 categories I-VIII (*14*). Genes belonging to categories I-IV and categories V-VII are activated and repressed by glucose, respectively (21). Genes in class VIII (N=804) are not regulated by glucose in wild type but only in *pka-as* cells and were excluded from further analysis. We compared the remaining list of glucose-responsive genes (N=2469) with the list of 2426 TORC1 target genes reported in a recent transcriptomic study (*14, 19*). We found that 58 % of glucose-responsive genes (N=1436; upregulated=828 and downregulated =608) were co-regulated by TORC1 (Figure 3C). Among the 828 genes upregulated by glucose, 807 (98%) genes were positively regulated by TORC1. Likewise, of the 608 genes negatively regulated by glucose, 578 (95%) genes were also negatively regulated by TORC1. Genes regulated by TORC1 were spread across the seven categories of glucose-responsive genes to different extents (Figure 3D). These observations indicate that addition of glucose and TORC1 activation have similar qualitative effects on the yeast transcriptome.

### TORC1 is required for establishment and maintenance of TGC gene expression

We investigated whether TORC1 is required for the transcriptional response to glucose. We chose 7 TORC1 target genes *GFD2* (I)*, DHR2* (II), *CIT1*(V), *CRC1*(V), *UGA1*(VI), *RME1*(VI) and *GPG1*(VII) spread across the top 5 expression categories as representative of ‘**T**ORC1 and **G**lucose **C**o-regulated (**TGC**)’ genes for analysis. The choice of these 7 genes was also informed by our transcriptome analysis of TORC1 targets (see below). We tested whether TORC1 activity is necessary for glucose-induced changes in expression of the seven TGC genes. We added glucose (to a final concentration of 2%) to log-phase yeast cells growing in SC/EG (SC medium containing 2% ethanol and 2% glycerol), in the presence and absence of rapamycin. Expression of all the 7 genes was significantly altered after 30 minutes following addition of glucose (Figure 4). As expected, *GFD2* and *DHR2* genes (belonging to clusters I and II respectively) were upregulated in the presence of glucose (Figure 4). The genes *GPG1*, *UGA1*, *RME1*, *CIT1* and *CRC1* (belonging to clusters V-VII) were downregulated in the presence of glucose (Figure 4). Importantly, the glucose-induced changes in expression of the 7 TGC genes were abolished by addition of rapamycin (Figure 4). These results indicate that TORC1 activity is required for the transcriptional response to glucose.

**Figure 4.**
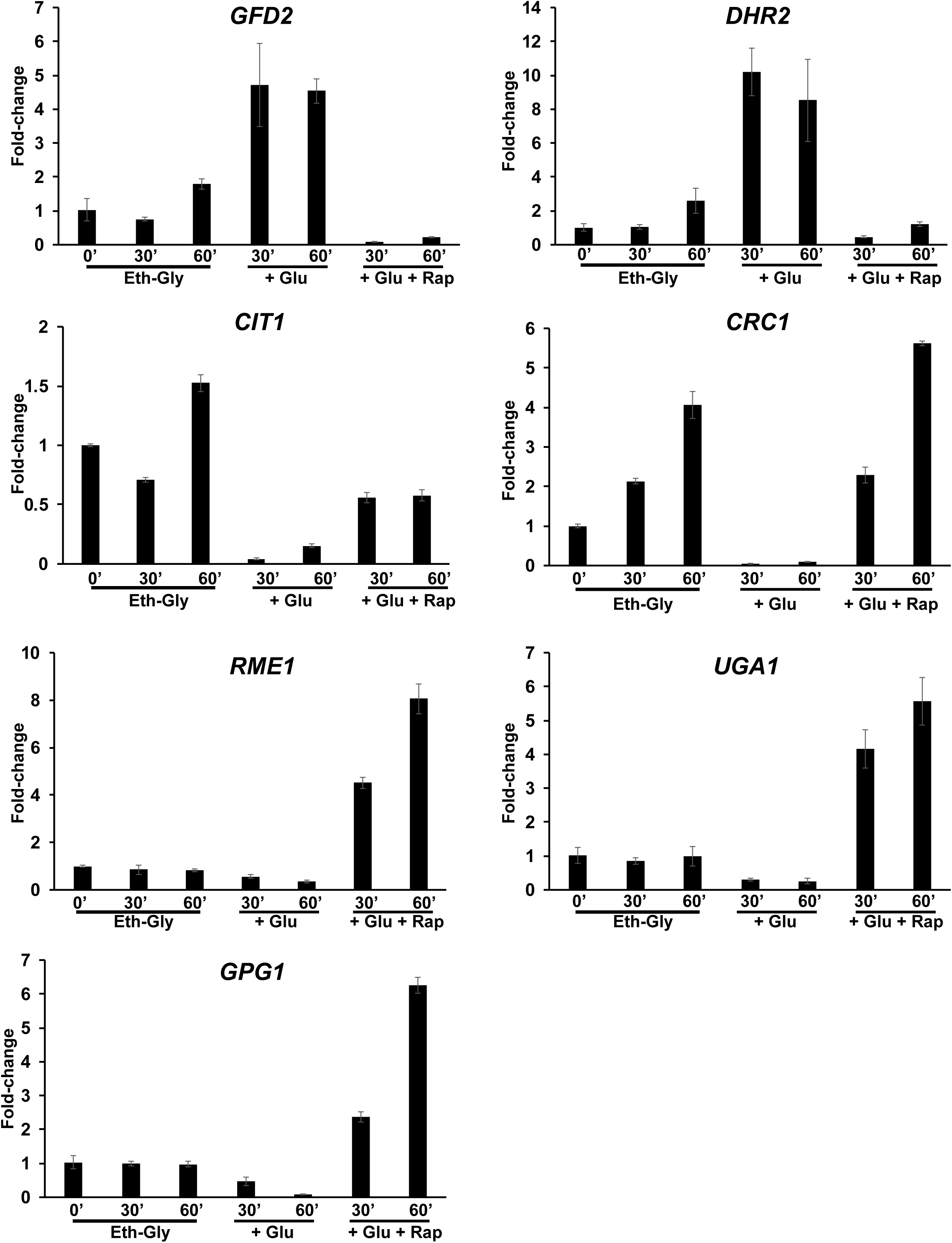
TORC1 is required for the transcriptional response to glucose. Wild type cells were grown to logarithmic phase in SC/EG medium and then glucose (2% final concentration) was added to the cultures in the presence of either rapamycin (200 nM) or DMSO. Aliquots of the cultures were taken after 30’ and 60’. RNA was extracted from the cultures and expression of 7 glucose-responsive genes (*GFD2*, *GPG1*, *UGA1*, *RME1*, *CIT1*, *CRC1* and *DHR2*) was assayed by Real-Time qRT-PCR analysis. Fold-change values were determined by normalizing with respect to cells in SC/EG medium at t=0’.

We then tested whether maintenance of TGC gene expression in glucose-containing medium is dependent on TORC1 activity. We treated log phase yeast cells growing in SC/D (SC medium containing glucose) with either DMSO or rapamycin (200 nM) and analyzed the expression of the 7 TGC genes by Real-Time qRT-PCR. Addition of rapamycin affected the expression of all the 7 genes with the transcript levels shifting towards the corresponding expression level in SC/EG medium (Figure S2). These results indicate that TORC1 activity is also required for maintaining the expression status of TGC genes during growth in glucose-containing medium (Figure S2).

### TORC1 and PKA co-regulate the expression of glucose-responsive genes

Glucose-responsive genes were reported to be primarily regulated by the PKA/Ras cAMP pathway in yeast (*2, 14*). However, another study suggested that TORC1 works with PKA in regulating the expression of glucose-responsive genes (*16*). We added glucose (2%) to wild type and *pka-as* strains growing in SC-EG medium along with either DMSO or 1-NM-PP1 or rapamycin and examined the levels of the 7 TGC genes by Real-Time qRT-PCR. As expected, glucose-induced changes in expression of all the seven genes were affected by rapamycin treatment in both PKA and *pka-as* strains (Figure 5). For two genes (*GFD2* and *DHR2*), 1-NM-PP1 had a non-specific effect on gene expression (Supplementary Figure S3). However, the expression of the remaining 5 TGC genes (*CIT1*, *CRC1*, *UGA1*, *RME1* and *GPG1*) was affected by 1-NM-PP1 specifically in the *pka-as* strain but not in the PKA strain (Figure 5 and Supplementary Figure S3). Our results indicate that the TGC genes are largely co-regulated by TORC1 and PKA.

**Figure 5.**
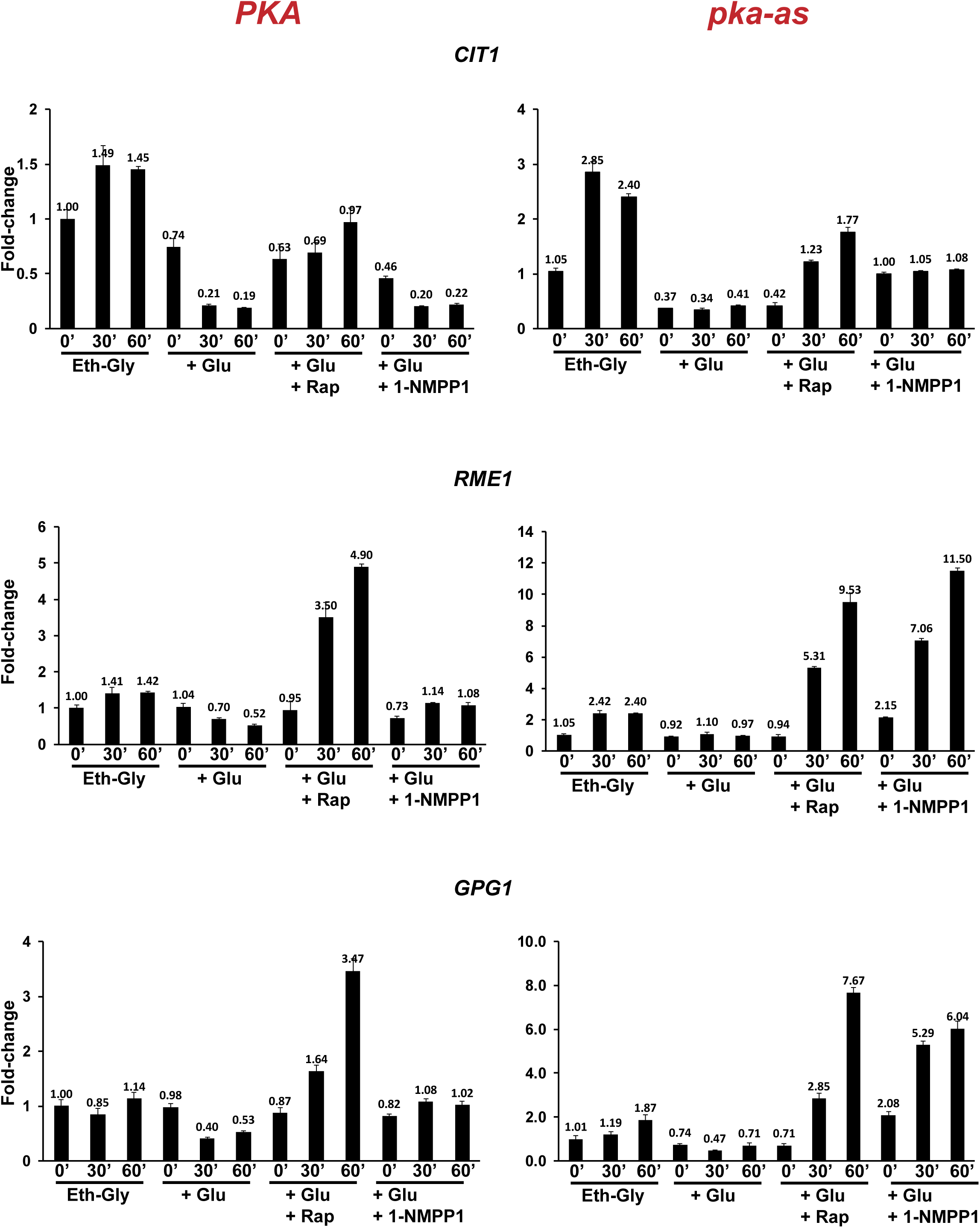
PKA and TORC1 co-regulate the glucose-responsive genes. Wild type (PKA) and *pka-as* cells were grown to logarithmic phase and then treated with either rapamycin (200 nM) or 1-NM-PP1 (25 µM) or DMSO. Aliquots of the cultures were taken after 0, 30’ and 60’. RNA was extracted from the cultures and the expression of the indicated 7 TGC genes were analyzed by Real-Time qRT-PCR. Results for *GPG1*, *RME1* and *CIT1* are presented here. Results for *GFD2*, *UGA1*, *CRC1* and *DHR2* are presented in Supplementary Figure S3.

### TORC1 regulates the expression of glucose-responsive genes independently of Bcy1 phosphorylation

TORC1 could regulate the expression of glucose-responsive genes indirectly by activating PKA. Indeed, TORC1 has been shown to promote PKA activity by inhibiting phosphorylation of PKA inhibitor Bcy1 at T129 (*20*). Phospho-mimetic mutation of T129 (*bcy1-T129D*) was shown to have an inhibitory effect on PKA activity (21). If TORC1’s effect on PKA activity *via* regulating T129 phosphorylation is important for the transcriptional response to glucose, then *bcy1-T129D* should block TORC1’s role in the glucose-response. We added glucose to *BCY1* and *bcy1-T129D* cultures growing in SC-EG medium and monitored the expression of three TGC genes *DHR2*, *CIT1* and *RME1* by Real-Time qRT-PCR. Rapamycin treatment affected the expression of the TGC genes to comparable extents in both the *BCY1* and *bcy1-T129D* strains (Supplementary Figure S4) indicating that TORC1 regulates the glucose-responsive genes independently of Bcy1 phosphorylation at T129.

### TORC1 inhibition does not affect PKA activity

TORC1 and PKA have also been shown to exert mutually antagonistic effects on their activities (*21*). We therefore tested the effect of TORC1 inhibition on *in vivo* PKA activity. We treated logarithmically growing wild type or *pka-as* cells with either 1-NM-PP1 or rapamycin or DMSO. We assayed PKA activity by western blotting of whole cell extracts using a phospho-specific antibody directed against phosphorylated PKA substrates (*20*). As expected, 1-NM-PP1 inhibited the phosphorylation of PKA substrates in the *pka-as* but not in the wild type strain (Supplementary Figure S5). In contrast, rapamycin treatment did not affect phosphorylation of PKA substrates (Supplementary Figure S5). However, rapamycin treatment affected the expression of TORC1 target genes *DIP5* and *GAP1* in *PKA* and *pka-as* cells (Supplementary Figure S5). Our data suggest that TORC1 inhibition does not affect PKA activity.

### TORC1 regulates the expression of glucose-responsive genes independently of Sch9

We then tested whether TORC1-mediated regulation of TGC genes is dependent on its downstream effectors Sch9 and Tap42. As PKA and TORC1 co-regulate the glucose-response, inactivating the TORC1 effector involved in glucose-response might not have a major effect on glucose-induced gene expression. However, rapamycin treatment will be expected to have a reduced effect on expression of glucose-responsive genes in the effector mutant strain in comparison to the wild type strain. To assess the role of Sch9 in TORC1-mediated regulation of the TGC genes, we added glucose (2% final) to wild type and *sch9Δ* cultures growing in SC/EG medium. We assayed the expression of the 7 TGC genes in the presence and absence of rapamycin by Real-Time qRT-PCR. Upregulation of *GFD2* and *RME1* after 30’ following addition of glucose was reduced in the *sch9Δ* strain in comparison to the wild type strain.

However, the expression of the remaining five downregulated TGC genes was reduced similarly in the wild type and the *sch9Δ* strains following addition of glucose (Figure 6). Importantly, addition of rapamycin reversed the glucose-induced changes in expression of all the 7 TGC genes in both wild type and *sch9Δ* strains. However, the effect of rapamycin on TGC gene expression was reduced in the *sch9Δ* strain in comparison to the wild type strain. Our data indicate that although Sch9 can boost the transcriptional response to glucose, TORC1 can regulate the glucose-responsive genes independently of Sch9 (Figure 6).

**Figure 6.**
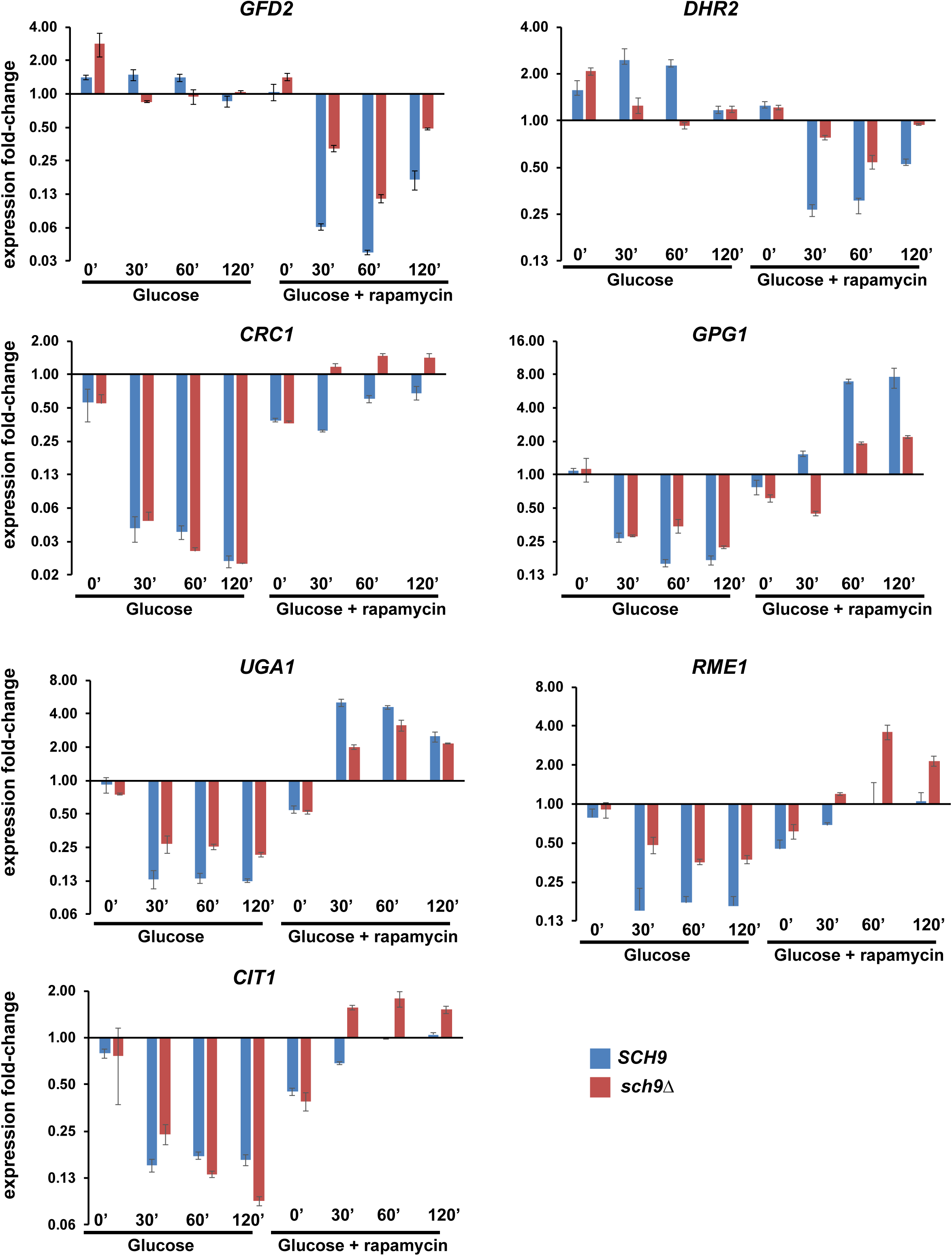
TORC1 regulates the expression of glucose-responsive genes independently of Sch9. Wild type and *sch9Δ* cells were grown to logarithmic phase in SC/EG medium and then glucose (2% final concentration) was added to the cultures in the presence of either rapamycin (200 nM) or DMSO. Aliquots of the cultures were taken after 0’, 30’, 60’ and 120’. RNA was extracted from the cultures and the expression of the indicated 7 glucose response genes (*GFD2*, *GPG1*, *UGA1*, *RME1*, *CIT1*, *CRC1* and *DHR2*) were analyzed by Real-Time qRT-PCR.

### TORC1 regulates the expression of glucose-responsive genes *via* Tap42

To test the role of Tap42 in the expression of glucose-responsive genes, we used *tap42-11* a temperature-sensitive allele of *TAP42* (*22*). We took wild type and *tap42-11* cells growing in mid-log phase at 25 °C (permissive temperature) in SC/EG medium and transferred them to 37 °C (non-permissive temperature) for about 30’ to inactivate Tap42. We then added 2% glucose to the cultures along with either rapamycin or DMSO and assayed the expression of the 7 TGC genes by Real-Time qRT-PCR. Induction of *GFD2* and *DHR2* expression after 30’ following addition of glucose was severely inhibited in *tap42-11* cells (Figure 7). Furthermore, addition of rapamycin had a reduced effect on *DHR2* expression in *tap42-11* cells in comparison to wild type cells suggesting that TORC1 regulates *DHR2* expression mainly *via* Tap42. For the remaining 5 TGC genes, addition of glucose affected their expression to comparable extents in both wild type and *tap42-ts* strains (Figure 7). However, interestingly addition of rapamycin had little or no effect on expression of *UGA1*, *RME1* and *GPG1* genes in *tap42-ts* cells (Figure 7). In addition, the effect of rapamycin on *CRC1* expression in *tap42-11* cells was reduced by 4-fold at 120’ in comparison to the wild type cells. These results indicate that TORC1 regulates glucose-driven changes in at least 4 (*DHR2*, *UGA1*, *RME1* and *GPG1*) of the 7 TGC genes through Tap42. It is possible that both Sch9 and Tap42 co-regulate the remaining 3 TGC genes. We were unable to test this possibility as *tap42-11* was synthetic lethal with *sch9*Δ.

**Figure 7.**
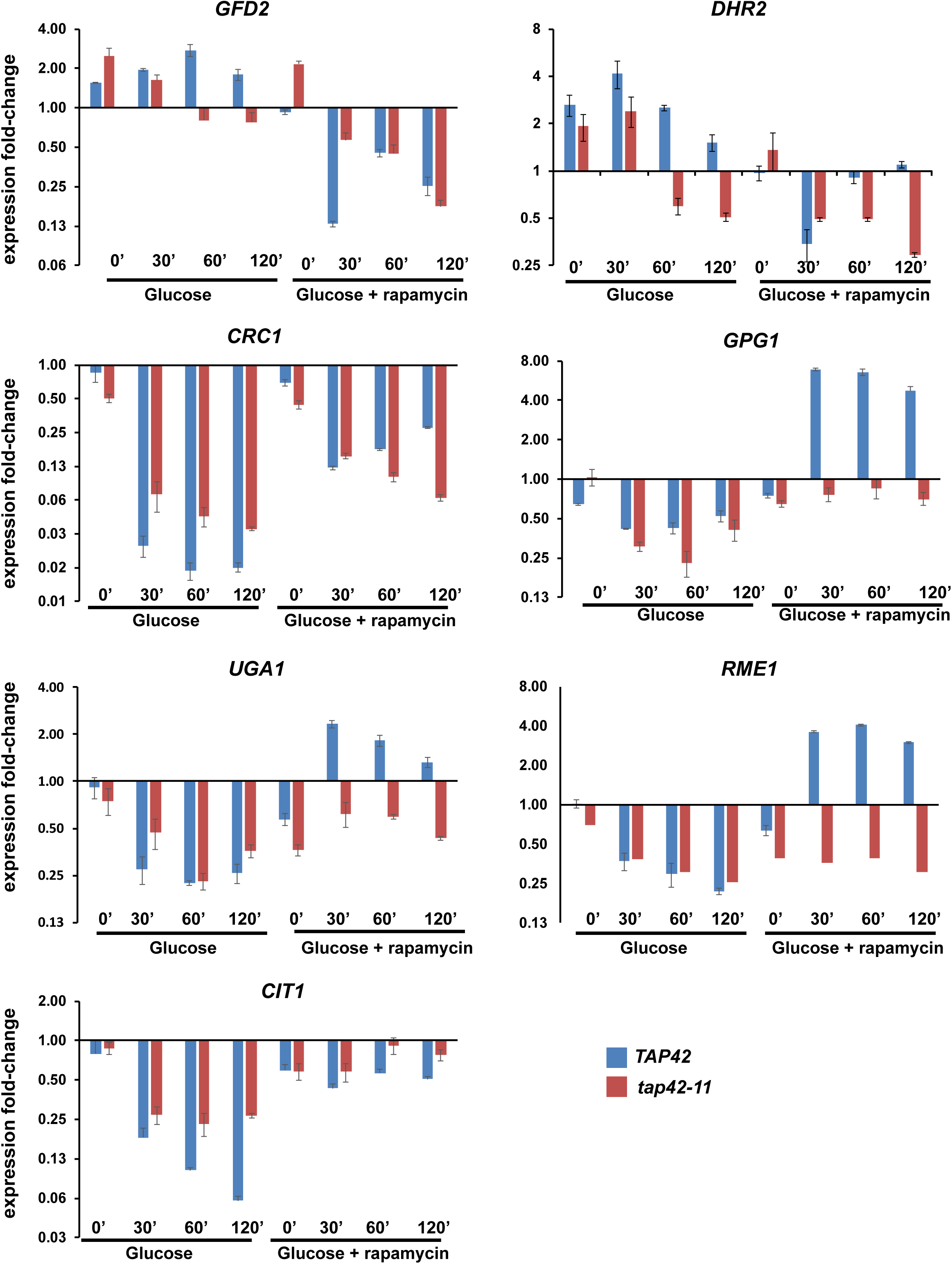
Regulation of glucose-responsive genes by TORC1 is dependent on Tap42. Wild type or *tap42-11* cells were grown to logarithmic phase at 25 °C (permissive temperature) in SC/EG medium and then shifted to 37°C for 30’ to inactivate *tap42-11*. Glucose (2% final concentration) was added to the cultures in the presence of either rapamycin (200 nM) or DMSO. in the presence or absence of rapamycin (200 nM). Aliquots of the cultures were taken after 0’, 30’, 60’ and 120’. RNA was extracted from the cultures and the expression of the 7 glucose response genes (*GFD2*, *GPG1*, *UGA1*, *RME1*, *CIT1*, *CRC1* and *DHR2*) were analyzed by Real-Time qRT-PCR.

### Rrd1/Rrd2 and Sit4 proteins are required for TORC1’s role in the transcriptional response to glucose

Tap42 interacts with the catalytic subunit of PP2A phosphatases (PP2Ac) and PP2A-like phosphatase (Sit4) in log phase but not in stationary phase cells and keeps them inactive and localized to the vacuole (*23*). Upon starvation (or rapamycin treatment), the phosphatases dissociate from Tap42 and dephosphorylate their target proteins in collaboration with two Phosphotyrosyl Phosphatase Activator (PTPA) proteins Rrd1 and Rrd2 (*24*). We tested whether the TORC1-regulation of TGC genes is dependent on PP2A-like phosphatase Sit4, Rrd1 and Rrd2. As *sit4Δ* and *rrd1Δ rrd2Δ* strains were unable to grow in the presence of glycerol as a carbon source, we tested their roles in the maintenance of TGC gene expression. We treated wild type, *sit4Δ*, *rrd1Δ, rrd2Δ* and *rrd1Δ rrd2Δ* cells growing in YPD/D (with 2 % glucose) medium with either DMSO or rapamycin and assessed the expression of TGC genes by Real-Time qRT-PCR. As expected, rapamycin treatment decreased the expression of *GFD2* and *DHR2* genes and increased the expression of *CIT1*, *CRC1*, *UGA1*, *RME1* and *GPG1* genes in wild type cells.

Rapamycin-induced changes in expression of TGC genes were reduced considerably in *rrd1Δ rrd2Δ* and *sit4Δ* cells (Figure 8 and Supplementary Figure S6). Rapamycin-induced changes in TGC gene expression were slightly reduced in the *rrd1*Δ strain but not in the *rrd2Δ* strain.

**Figure 8.**
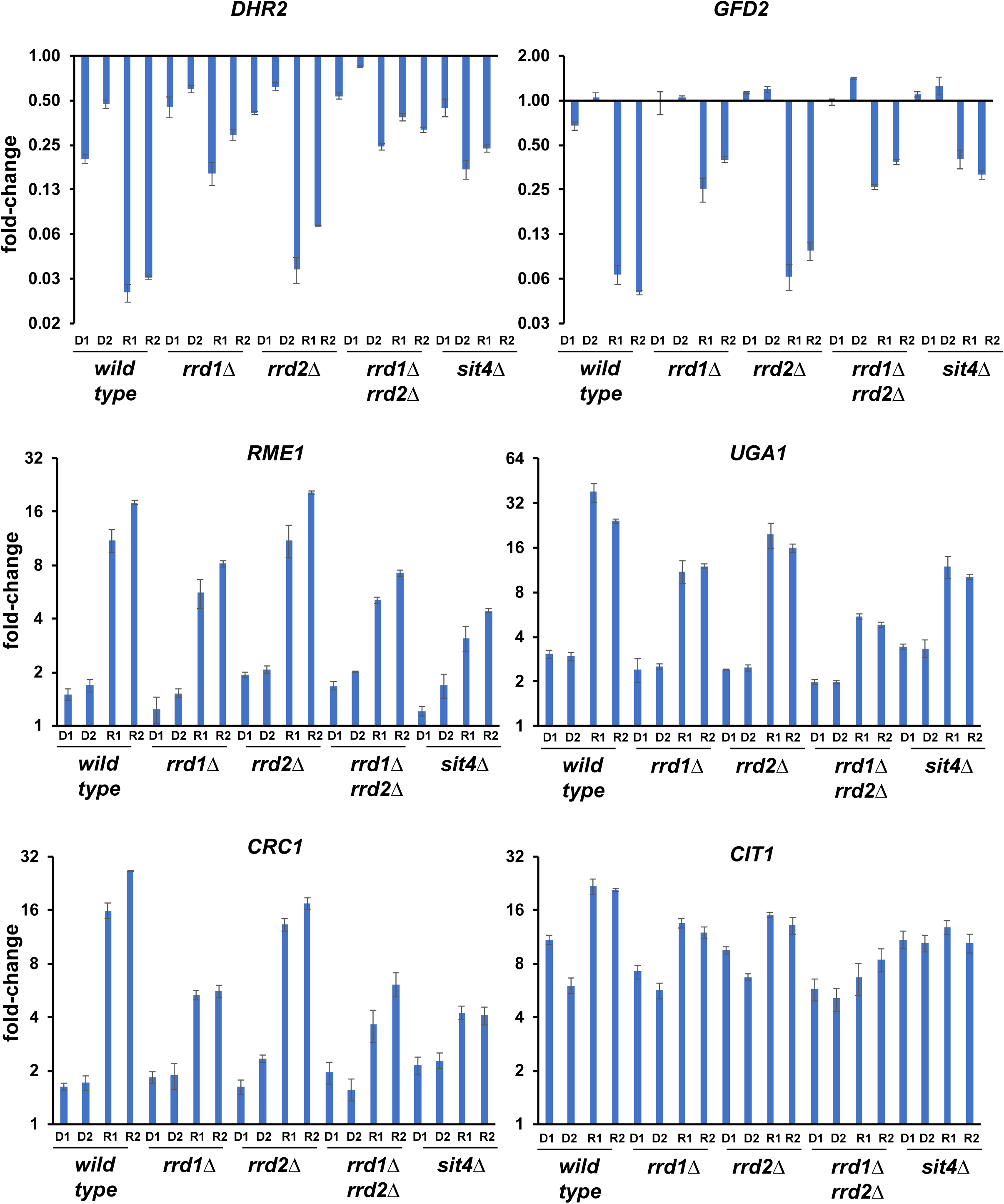
TORC1 regulates the glucose-responsive genes via Sit4 and Rrd1/Rrd2 proteins. Wild type, *sit4Δ, rrd1Δ, rrd2Δ* and *rrd1Δ rrd2Δ* cells were grown to logarithmic phase and then treated with either rapamycin (200 nM) or DMSO. Aliquots of the cultures were taken after 0’, 30’ and 60’. RNA was extracted from the cultures and the expression of the indicated 7 TGC genes was analyzed by Real-Time qRT-PCR. D1/R1 and D2/R2 indicate DMSO-treated/rapamycin-treated cells after 30’ and 60’ respectively and the expression fold-change values were normalized with respect to DMSO-treated cells at t=0’. Results for *GFD2*, *UGA1*, *RME1*, *CIT1*, *CRC1* and *DHR2* are presented here. Results for *GPG1* are in Supplementary Figure S6.

However, the rapamycin-induced changes in the *rrd1Δ rrd2Δ* strain were less than in the *rrd1Δ* strain (Figure 8 and Supplementary Figure S6) suggesting that Rrd1 and Rrd2 proteins play an overlapping role in TORC1-mediated regulation of TGC genes. Taken together, our results are consistent with the hypothesis that TORC1 regulates the glucose-response genes by inhibiting the activities of PP2A/Sit4/Rrd1/Rrd2 phosphatases *via* Tap42.

### Glucose activates TORC1 in spores

To test the physiological importance of TORC1’s role in glucose-signaling, we investigated the function of TORC1 in spore germination. Return of spores into vegetative growth cycle upon their transfer to favorable nutrient medium is referred to as spore germination. Glucose (or a fermentable carbon source) is essential for efficient spore germination (*25*). Glucose alone is sufficient to trigger spore germination (*25*). Moreover, spores are only responsive to glucose during the early stages of germination. Consistent with this, the transcriptional changes induced upon transfer of spores into rich medium and glucose are strikingly similar during the initial stages of spore germination (*1, 26*).

We first tested whether TORC1 is activated upon transfer of spores into nutrient medium. We generated spores by transferring stationary phase wild type diploid cells expressing HA-tagged Sch9 pre-grown in nutrient medium (YPD) for 16-20 hours, into sporulation medium. After 84 hours of incubation at 30 °C in sporulation medium, more than 90% of diploid cells had sporulated (data not shown). We purified the spores and transferred them into nutrient medium (YPD) in the presence of rapamycin (2 µM) or DMSO (mock-treated). Sch9 was completely dephosphorylated in spores indicating that TORC1 is inactive in the spore (Figure 9A).

**Figure 9.**
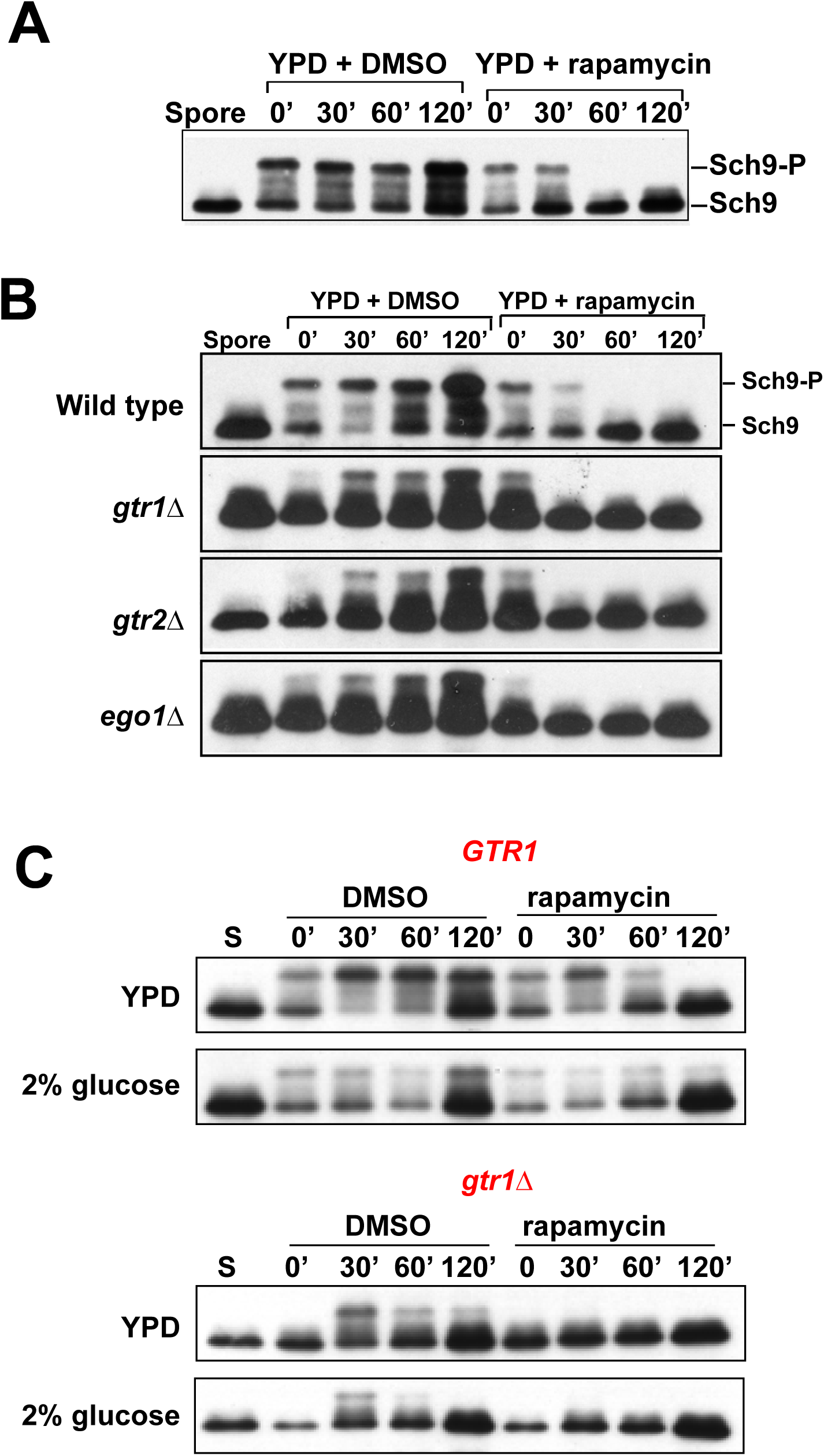
Glucose activates TORC1 in spores by Gtr1/2-dependent and Gtr1/2-independent mechanisms. A) Wild type spores expressing Sch9-HA6 were transferred into nutrient medium in the presence of either DMSO or rapamycin (2 µM). Activation of TORC1 was assayed by examining phosphorylation of Sch9 by Western blotting using an anti-HA antibody. B) Wild type spores and spores bearing *gtr1Δ* / *gtr2Δ* / *ego1Δ* were transferred into nutrient medium in the presence of either DMSO or rapamycin (2 µM). Activity of TORC1 was assayed by monitoring Sch9 phosphorylation using an anti-HA antibody. C) Wild type spores and *gtr1Δ* spores were transferred into either nutrient medium or glucose (110 mM) in the presence of either DMSO or rapamycin (2 µM). Activity of TORC1 was assayed by monitoring Sch9 phosphorylation using an anti-HA antibody.

Immediately after transfer of spores into nutrient medium, Sch9 was phosphorylated (Figure 9A). Addition of rapamycin to the nutrient medium inhibited Sch9 phosphorylation after 60’ (Figure 9A) indicating that TORC1 is activated immediately following transfer of spores into nutrient medium.

To test whether Gtr1/ Gtr2 and EGO complexes are required for TORC1 activation during spore germination, we generated wild type, *gtr1Δ*, *gtr2Δ* and *ego1Δ* spores and transferred them onto nutrient medium and assayed TORC1 activation by monitoring Sch9 phosphorylation. Activation of TORC1 was reduced in *gtr1Δ*, *gtr2Δ* and *ego1Δ* spores in comparison to wild type spores (Figure 9B). In addition, rapamycin treatment of *gtr1Δ*, *gtr2Δ* and *ego1Δ* spores led to complete loss of Sch9 phosphorylation after 30’, in comparison to 60’ for the wild type strain following transfer into nutrient medium. Our data indicate that TORC1 activation is mediated by both EGO/Gtr1-Gtr2-dependent and EGO/Gtr1-Gtr2 -independent mechanisms during spore germination.

We then investigated whether glucose is sufficient to activate TORC1 in spores as noted in mitotically growing cells. We transferred wild type and *gtr1*Δ spores into either nutrient medium or glucose (2%). Incubation of wild type spores with nutrient medium and glucose resulted in immediate rapamycin-sensitive Sch9 phosphorylation (Figure 9C). Activation of TORC1 by nutrient medium and glucose was weaker and delayed in *gtr1Δ* spores. These results indicate that glucose is sufficient to activate TORC1 *via* Gtr1/2-dependent and -independent mechanisms in spores (Figure 9C).

### Transcriptomic analysis of spore germination

To test whether TORC1 regulates the glucose-responsive genes during spore germination, we performed transcriptomic analysis of germinating spores transferred to nutrient medium in the presence and absence of rapamycin. In addition to purified spores, we collected germinating yeast spores at different time points (0’, 10’, 30’, 60’, 120’, 240’ and 360’) following their transfer into nutrient medium (in the presence and absence of rapamycin) and analyzed their transcriptomes by RNA-Seq (Figure 10A).

**Figure 10.**
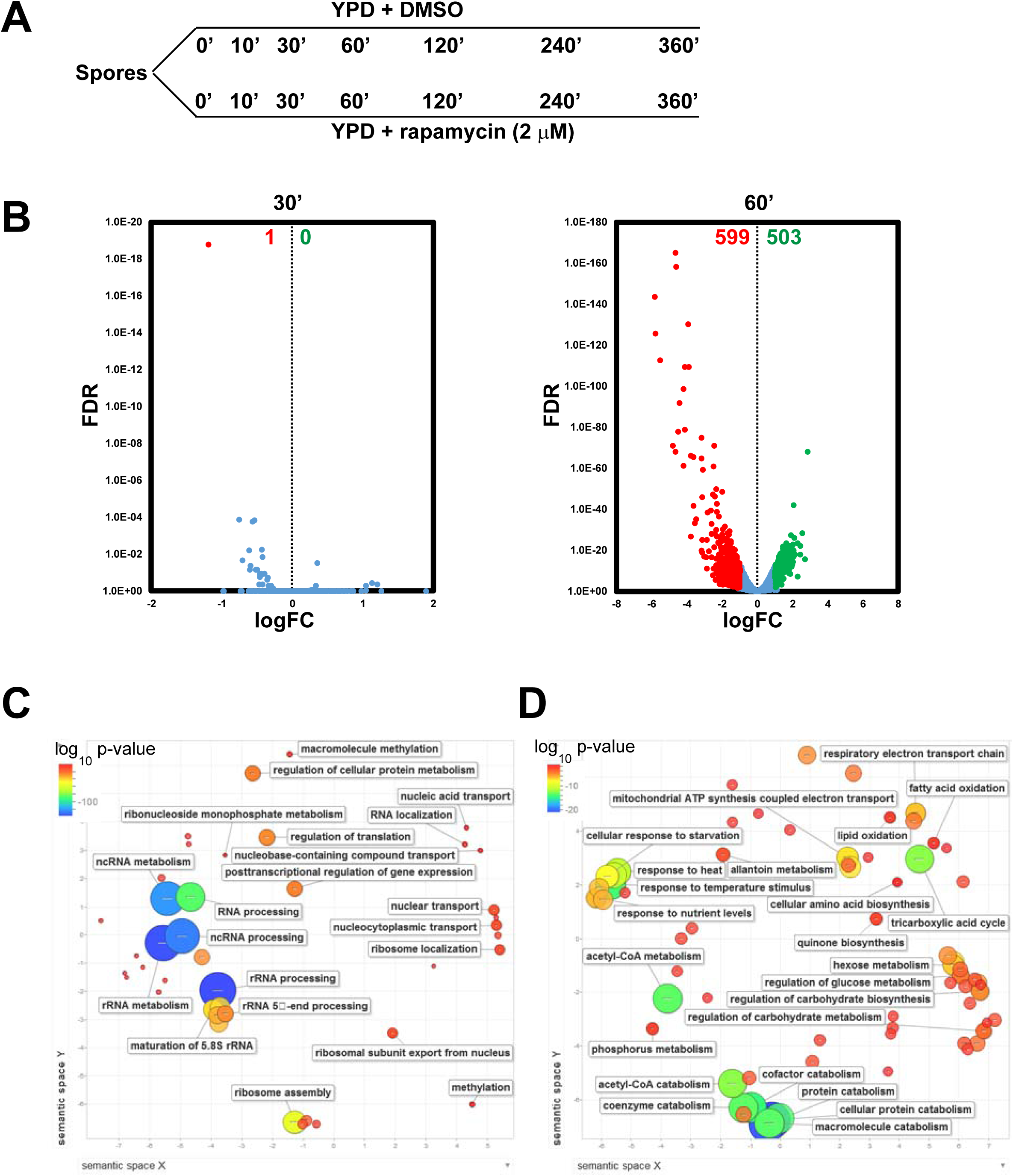
TORC1 regulates transcriptomic changes during spore germination. A) Experimental outline to analyze the role of TORC1 in spore germination by RNA-Seq. Spores were transferred to YPD medium containing either DMSO or rapamycin (2 µM). Aliquots of yeast cells were taken at the indicated time points (0’ 10’, 30’, 60’, 120’, 240’ and 360’) from the two cultures and used for preparing RNA for RNA-Seq analysis. B) Transcriptomes of spores treated with rapamycin (2 µM) after 30’ and 60’ was compared with the transcriptomes of corresponding DMSO-treated spores. logFC (Fold Change) was plotted against False Discovery Rate (FDR). Differentially Expressed Genes (at least 2-fold difference in comparison to DMSO-treated spores) were identified. Genes positively regulated and negatively regulated by TORC1 are indicated by green and red dots respectively and their numbers are indicated at the top of the plot. C) Representation of enriched GO terms among genes that are upregulated by TORC1 (at least 2-fold increase) during spore germination by REVIGO. D) Same as C but for TORC1-downregulated genes (at least 2-fold decrease).

To examine changes in the transcriptome during spore germination, we compared the transcript levels of genes in germinating spores (YPD + DMSO) with the corresponding transcript levels in the ungerminated spore and identified differentially expressed genes that showed at least a 2-fold difference at the 7 timepoints (data file S1). The transcriptome of spores transferred to YPD for t=0’ matched well with that of the spores with only 24 genes being differentially expressed (Supplementary Figure S7 and Supplementary table S1). The number of differentially expressed genes increased with time and after 240’, 3458 genes were either upregulated (1733) or downregulated (1725) (Supplementary Figure S7 and Supplementary Table S1). Our RNA-Seq results are largely consistent with previous transcriptomic analyses of spore germination (*1, 26*). Ten classes of genes namely Protein synthesis, rRNA processing, Gluconeogenesis, TCA sub-cycle, Stress, Oxidative phosphorylation, Proteasome subunits, Mating and Cell cycle G1 and Cell cycle G2/M were reported to be differentially expressed during spore germination (*1*). We found that the aforementioned 10 classes of genes were also differentially expressed following the onset of germination in our experiment (Supplementary Figure S8).

### TORC1 regulates the glucose-responsive genes during spore germination

To determine how TORC1 regulates the transcriptome during spore germination, we compared the transcriptomes of DMSO- and rapamycin-treated spore germination cultures (Supplementary Table S2). There was no significant difference between the transcriptomes at t=30’ between the two cultures (Figure 10B and Supplementary Table S2). However, a significant difference in gene expression between the two cultures was observed after 1 hour following transfer of spores into rich medium (Figure 10B). Around 1102 genes (503 upregulated and 599 downregulated) were found to be differentially expressed (by at least 2-fold) between the two cultures (Figure 10B and Supplementary Table S2). This timing is consistent with our observation that TORC1 activity as measured by Sch9 phosphorylation is inhibited by rapamycin after 1 hour following transfer of spores into nutrient medium (Figure 10B). To validate the RNA-seq data, we first compared the expression of TORC1 targets like *DIP5 (*Dicarboxylic amino acid permease) and *GAP1* (General Amino acid permease) in DMSO- and rapamycin-treated spores after 0’, 30’, 60’ and 120’ following transfer to nutrient medium. Expression of TORC1-downregulated genes *DIP5* and *GAP1* was enhanced by rapamycin treatment after 1 hour following transfer of spores to nutrient medium consistent with RNA-Seq data (Supplementary Figure S9).

We performed Gene Ontology analyses to assess enhancement of specific functionally related gene groups within the 1102 TORC1-regulated genes. We used REVIGO to remove redundant GO terms and to visualize the enriched terms (*27*). As expected, the TORC1 upregulated target genes are involved in translation such as rRNA processing, ribosome assembly, ribosome localization and regulation of translation (Figure 10C). In addition, the downregulated TORC1 target genes involved in stress response (includes macroautophagy genes), protein catabolism, allantoin metabolism (Nitrogen Discrimination Pathway) and TCA cycle were identified in our analysis (Figure 10D).

We compared our list of 1102 TORC1 target genes with the list of TORC1 targets obtained from a microarray analysis-based study (*28*) and a recent RNA-Seq study (*19*) both performed with vegetative cells. Only 90 (18%) of the TORC1 targets identified in the microarray-based study (*28*) were in our set (Figure S10). However, about 935 (84.8%) of the TORC1 targets identified by RNA-Seq (*19*) were in our target list (Supplementary Figure S10). We don’t know the precise reason for differences in the TORC1 targets predicted by microarray and RNA-Seq methods but the differences in sensitivities of the two transcriptomic approaches (*29*) could be a contributing factor.

### TORC1 is required for expression of glucose-responsive genes during spore germination

To test whether TORC1 regulates the glucose-responsive genes during spore germination, we compared our list of TORC1 targets during spore germination with the list of glucose-responsive genes identified during vegetative growth (*14*). About 71 % (791 /1102) of TORC1 targets were also glucose-responsive (Supplementary Figure S10). To validate this observation, we compared the expression of 7 TGC genes in DMSO- and rapamycin-treated germinating spore cultures after 0’, 30’, 60’ and 120’ following transfer to nutrient medium. *DHR2* and *GFD2* genes were upregulated following spore germination and the genes *CIT1, GPG1, RME1, CRC1* and *UGA1* were downregulated following transfer of spores into nutrient medium. These changes in gene expression were similar to what we observed when we added glucose to yeast cells growing in medium containing ethanol and glycerol. Importantly, the changes in expression of 7 TGC genes were inhibited by addition of rapamycin (Supplementary Figure S11). These results indicate that TORC1 regulates the glucose-responsive genes during spore germination.

### TORC1 is essential for spore germination

Glucose is essential for efficient germination of yeast spores (*25*). If glucose-signaling is required for spore germination, then blocking either TORC1 or PKA should inhibit spore germination. We transferred wild type spores into nutrient medium in the presence of either rapamycin (2 µM) or DMSO (mock-treated) and followed the kinetics of spore germination by bright-field microscopy (Figure 11 A and B). After 2-3 h following transfer into nutrient medium, spores increased in volume and adopted a pear-shaped appearance (Figure 11 A and B). The spores elongated further and displayed a distinct constriction after 4 hours. The bud emerged from the mother cell subsequently and detached from the mother cell resulting in the first mitotic division after about 7-8 hours following transfer to nutrient medium (Figure 11A and B). However, in the presence of rapamycin, spore germination was severely inhibited. After 6 to 8 hours following rapamycin treatment, 50 % of spores had become enlarged but they did not form a constriction seen in untreated cultures and failed to undergo the first mitotic division (Figure 11 A and B). To test whether the effect of rapamycin on spore germination is due to inhibition of the TORC1 complex, we introduced the *TOR1-1* mutation that confers resistance to rapamycin (*18*). Rapamycin had no effect on germination of *TOR1-1* spores indicating that the effect of rapamycin on spore germination was due to specific inhibition of TORC1 (Figure 11B). In contrast to wild type spores, both *fpr1Δ* and *TOR1-1* spores germinated efficiently in the presence of rapamycin (Supplementary Figure S12).

**Figure 11.**
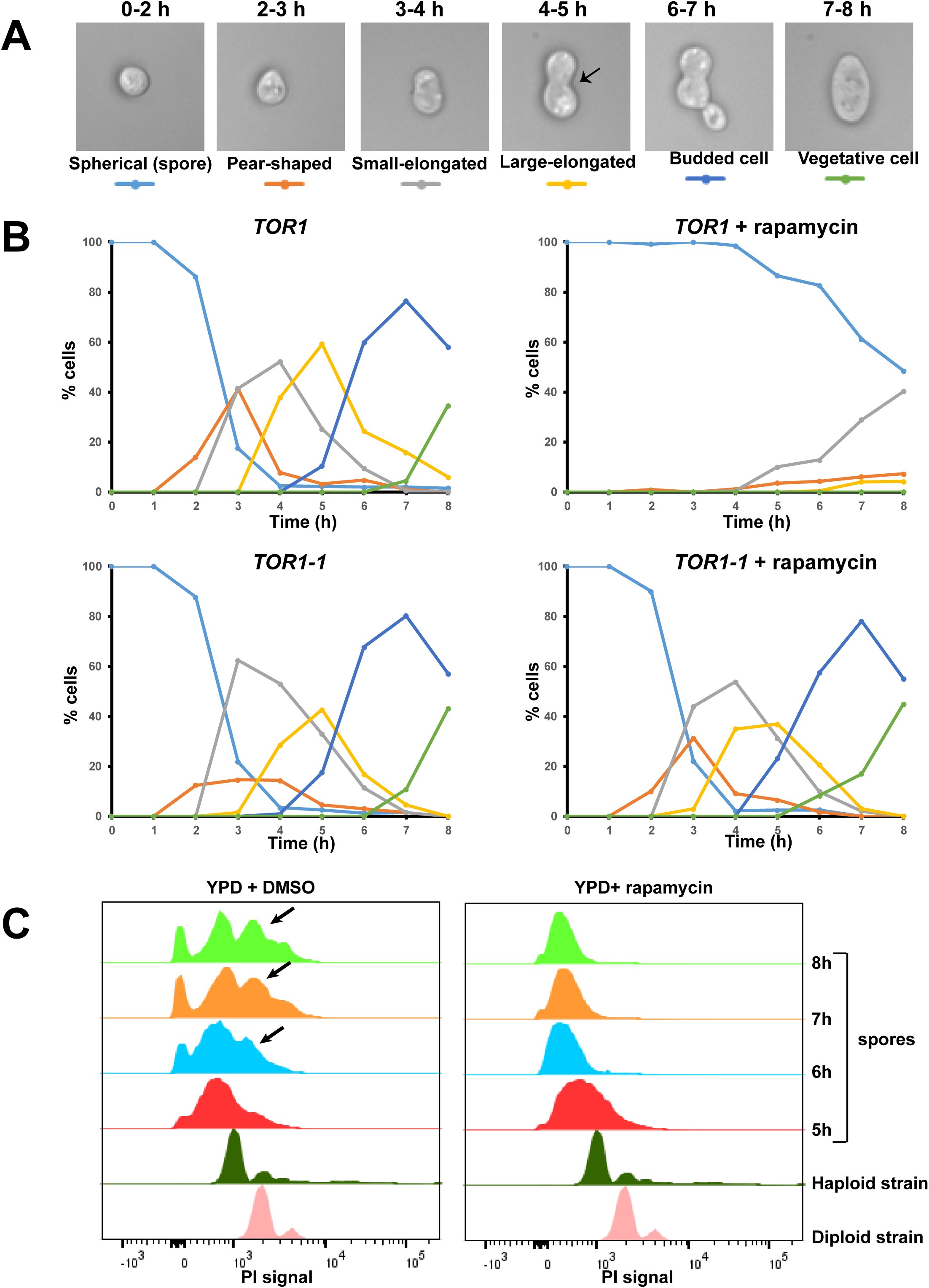
TORC1 activity is required for spore germination. A) Images showing the morphological changes occurring during spore germination along with their corresponding timing of appearance above. A distinct constriction seen in the germinating spores after 4-5 h following transfer to YPD medium is indicated by the black arrowhead. B) Wild type and *TOR1-1* spores were transferred into YPD medium in the presence of either DMSO or rapamycin (2 µM). Progress of spore germination was assayed by scoring fraction of cells with different morphologies at the indicated time points for 8 hours following transfer into YPD medium. C) Wild type spores were transferred into nutrient medium in the presence of either DMSO or rapamycin (2 µM). DNA content of germinating spores following 5 h, 6 h, 7 h and 8 h following transfer to YPD medium was assayed by flow cytometry. Propidium iodide signals of a haploid and diploid strain are indicated for reference.

We tested whether TORC1 is required for DNA replication, an event that occurs during final stages of spore germination (*1*). We transferred purified wild type spores into nutrient medium in presence and absence of rapamycin and monitored DNA replication by flow cytometry. The intensity of Propidium Iodide (PI) staining of spores was lower than that of mitotically growing haploid cells presumably because of reduced permeability to PI caused by the presence of spore wall. However, the wild type spores replicated their DNA between 6 and 8 hours as indicated by the appearance of cells with increased PI signals. In contrast, the PI signals of rapamycin-treated spores remained unchanged during the course of the experiment indicating that they did not undergo DNA replication (Figure 11C). We also confirmed that the expression of G1 cell cycle genes (*CLN1*, *CLN2*, *CLB6*, *EGT2*, *PCL1*, *PCL9*, *CHS2* and *CTS1*) expressed after 4 hours of germination was inhibited by rapamycin (Supplementary Figure S13). Taken together, our data indicate that TORC1 is required for spore germination in budding yeast.

We also tested whether Gtr1/Gtr2 complex is required for spore germination. Spores produced by *GTR1*/*GTR1* (wild type) and *gtr1Δ*/*gtr1Δ* diploid cells were purified and allowed to germinate by transferring them into nutrient medium. Wild type spores underwent the morphological changes as described above and 60% of spores were converted to budded cells after 7 hours following transfer to YPD medium (Supplementary Figure S14). As expected, addition of rapamycin to wild type spores prevented the formation of budded cells. The *gtr1Δ* spores germinated poorly with delayed kinetics in comparison to wild type spores (Supplementary Figure S14). While rapamycin-treated *GTR1* spores become elongated after 7 to 8 hours, rapamycin-treated *gtr1Δ* spores did not show any morphological change following transfer into nutrient medium (Supplementary Figure S14) and remained so even after 24 hours (data not shown). This suggests that the TORC1 is essential for all morphological changes that occur during spore germination and elongation of wild type spores observed in the presence of rapamycin (Figure 11A-B and Supplementary Figure S14) is due to incomplete inactivation of TORC1 by rapamycin. Consistent with this possibility, TORC1 activity is completely abolished by rapamycin treatment in *gtr1Δ* spores in comparison to wild type spores (Figure 9C).

We also tested whether PKA is essential for spore germination. We transferred wild type (*PKA*) and *pka-as* spores into nutrient medium in the presence of DMSO and 1-NM-PP1 and followed the kinetics of spore germination by bright-field microscopy. Addition of 1-NM-PP1 inhibited spore germination of *pka-as* spores but not of PKA spores (Supplementary Figure S15) indicating that PKA is essential for spore germination. This is consistent with an earlier report that spores from strains bearing temperature-sensitive alleles of Ras pathway fail to germinate at the non-permissive temperature (*25*).

### Effect of TORC1 inhibition on spore germination is dependent on Rrd1

If TORC1’s role in regulating the glucose responsive genes is essential for spore germination, then inactivating the Tap42/Sit4/Rrd1/Rrd2 branch should make spores germinate even in the presence of rapamycin. As *tap42-11*, *rrd1Δ rrd2Δ* and *sit4Δ* mutants failed to sporulate, we tested the effect of rapamycin treatment on germination of *rrd1Δ* spores. We took wild type and *rrd1Δ* spores and transferred them to nutrient medium in the presence and absence of rapamycin. Interestingly *rrd1Δ* spores germinated earlier in comparison to wild type spores as indicated by the formation of budded cells (Figure 12A). While the germination of wild type spores was inhibited by rapamycin treatment, the *rrd1Δ* mutant spores were able to form budded cells in the presence of rapamycin (Figure 12A). Our results demonstrate that the TORC1’s role in the regulation of glucose-responsive genes is essential for spore germination.

**Figure 12.**
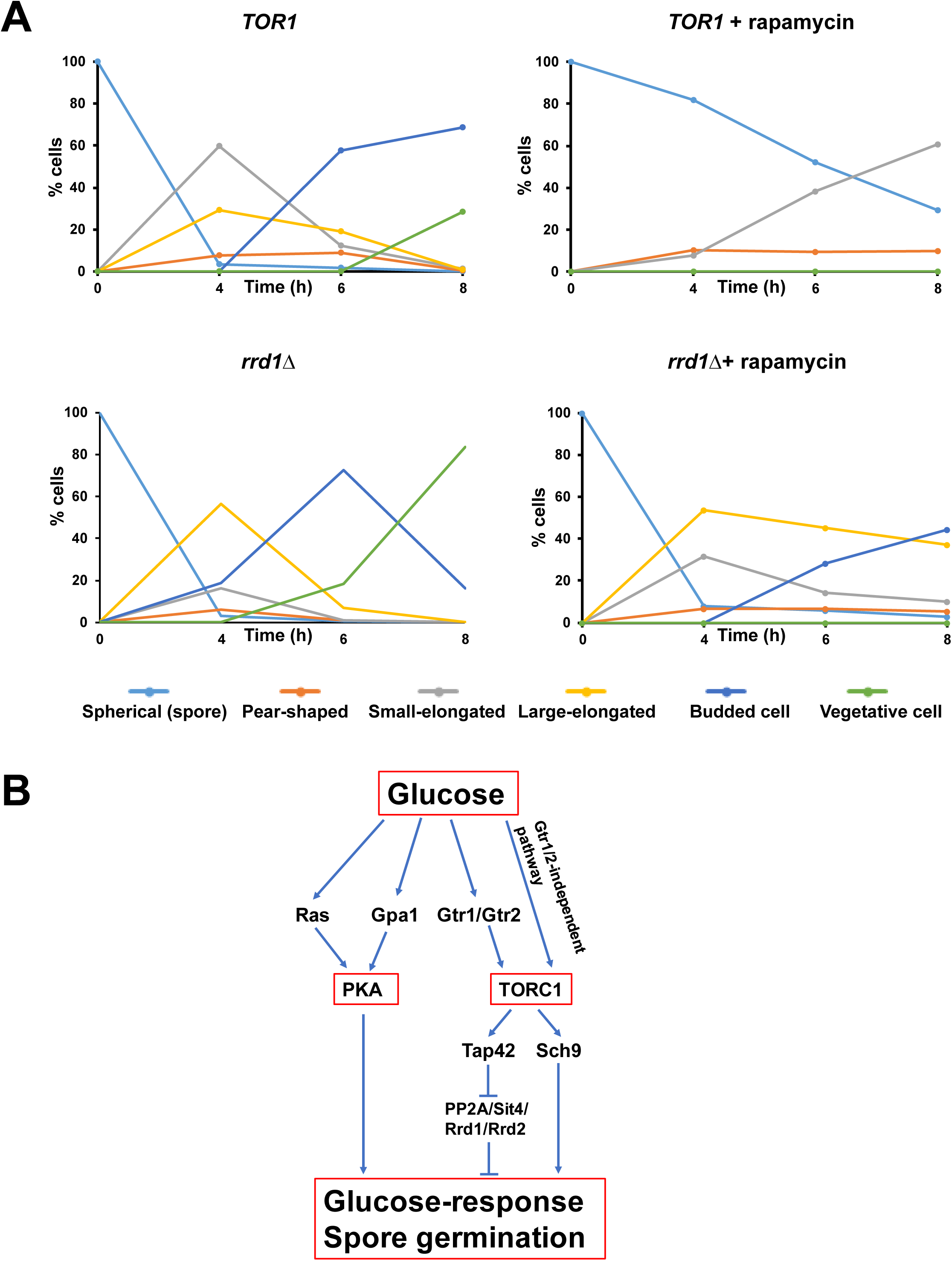
TORC1’s role in spore germination is mediated *via* inhibition of Rrd1. A) Wild type and *rrd1Δ* spores were transferred into YPD medium in the presence of either DMSO or rapamycin (2 µM). Progress of spore germination was assayed by scoring fraction of cells with different morphologies at the indicated time points for up to 8 hours following transfer into YPD medium. B) A model for how TORC1 regulates the glucose-response and spore germination via Tap42-Sit4-Rrd1/2 pathway. See the discussion for details.

## DISCUSSION

Target of Rapamycin Complex connects the presence of nutrients or growth factors in the environment with cellular growth and proliferation in eukaryotes. Dysregulation of TORC1 in humans has been associated with cancer, diabetes and obesity (*30*). Although TORC1’s function in coupling amino acid levels with cellular growth has been studied to considerable extent, its role in glucose-signaling is poorly understood and even somewhat controversial. Our work has enhanced the understanding of the relationship between glucose and TORC1 signaling in yeast. Firstly, we report that glucose is necessary and sufficient to activate TORC1. Transferring yeast cells from a medium containing a non-fermentable carbon source to glucose without changing the Nitrogen source causes robust activation of TORC1.

Secondly, TORC1 is required for the transcriptional response to glucose and this is accomplished mainly by inhibition of PP2A/PP2A-like phosphatases via its downstream effector Tap42. Finally, we have uncovered the physiological significance of TORC1’s role in glucose response by showing that TORC1 is required for spore germination, a glucose-dependent developmental transition in yeast.

Thus far, PKA and TORC1 have been implicated in sensing glucose and Nitrogen/amino acids levels respectively in yeast. We propose that TORC1 works with PKA in integrating information from glucose levels with growth and proliferation. A cellular nutrient sensor, in principle, should be activated by the nutrient and should prepare the cell for metabolism of the nutrient. By these two criteria, our study strongly suggests that TORC1 is a glucose-sensor in yeast (Figure 12B). Glucose activates TORC1 and TORC1 regulates the glucose-responsive genes (Figure 12B).

A recent study evaluated the contributions of PKA and TORC1 to transcriptional regulation of ribosomal biogenesis and protein synthesis genes following transfer of cells from a non-fermentable carbon source to glucose using chemical inhibitors (*31*). They found that while PKA was required for the regulation of ribosomal biogenesis/ protein synthesis genes during transition into glucose-containing medium, TORC1 was required for their steady state gene expression (*31*). Although they proposed that TORC1 regulates the ribosomal biogenesis genes by inactivating transcriptional repressors Dot6 and Tod6 via Sch9, rapamycin treatment affected the gene expression in a *dot6Δ tod6Δ* strain (*31*). This is consistent with our observations that rapamycin treatment affected the expression of the TGC genes in the *sch9Δ* strain (Figure 6). In contrast, rapamycin treatment had no/ significantly reduced effect on expression of glucose-response genes in strains lacking functional Tap42 or Sit4 or Rrd1/Rrd2 proteins in our experiments. Precisely how PP2A and PP2A-like phosphatase Sit4 regulate the transcription of glucose-response genes remains a topic that merits further investigation.

Another key question emerging from our study is how does glucose activate TORC1. Glucose deprivation has been reported to cause TORC1 inactivation (*17, 32, 33*). However, we show that glucose is also sufficient for TORC1 activation following complete nutrient starvation. This activation is partly dependent on Gtr1/Gtr2 complex but is independent of PKA activity. Snf1 kinase, which is active during glucose starvation, has been shown to inactivate TORC1 by phosphorylating its subunit Kog1 (*32*). In addition, glucose has also been reported to regulate TORC1 activity by altering the cytosolic pH which promotes a physical interaction between v-ATPase and Gtr1 (*34*). It will be interesting to test whether glucose-induced activation of TORC1 observed in our assay occurs *via* inhibition of Snf1 and/or activation of v-ATPase.

Several intriguing questions emerge from our study. Does glucose need to be metabolized by the cell to activate TORC1? Do the mechanisms of TORC1 activation by glucose and amino acids *via* the Gtr1/Gtr2 complex overlap? What factors regulate the Gtr1-independent pathway of glucose-mediated TORC1 activation? Interestingly in plants, glucose has been shown to activate TORC1 and the TORC1-glucose signaling is essential for transcriptional reprogramming and development (*35*). It is quite plausible that the TORC1-glucose signaling is an ancient conserved pathway employed by eukaryotic cells to coordinate presence of nutrients in the environment with their growth and developmental state.

## MATERIALS AND METHODS

### Yeast strains and plasmids

All strains are derived from SK1 genetic background. A complete list of yeast strains along with their genotypes can be found in Supplementary Table S3.

### TORC1 activity assay

Protein extraction, chemical fragmentation and western analysis were performed as previously described with slight modifications (*17*). Germinating spore culture collected at different time points was mixed with tricholoroacetic acid to a final concentration of 6%. Samples were then kept on ice for at least 5 min and further washed twice with cold acetone, and air dried in a fume hood. Lysis of the cells was done in 150 µl of urea buffer (50 mM Tris pH 7.5, 5 mM EDTA, 6 M urea, 1% SDS, 1 mM PMSF, and 1x PPi) with glass beads in a bead beater followed by incubation for 10 min to 65 °C in a thermomixer with shaking at 800 rpm. Then after protein either stored at -80 °C or performed NTCB cleavage assay. For NTCB cleavage 30 µl of 0.5 M CHES (pH 10.5) and 20 µl of NTCB (7.5 mM in H2O) were added and samples incubated over night at RT with shaking. Then after 1 vol of 2x sample buffer (2x SDS gel-loading buffer, 20 mM TCEP and 0.5x PPi) was added to the samples. Further proteins were separated on 8% SDS-PAGE gels and transferred onto nitrocellulose membrane. Blots were then blocked with 5% milk in PBS/0.1% Tween 20. Blots were probed with anti-HA antibody 16B12 (1:1000; cat no. 901501, BioLegend) or 3F10 (1:1000; cat no. 11867431001, Roche) followed by sheep anti-mouse antibody (1:5000; ECL^TM^) or goat anti-rat antibody (1:5000; Santa-Cruz Biotechnology) conjugated with horseradish peroxidase. Finally, the blots were developed with ECL prime western blotting detection reagent (Amersham Pharmacia Biotech).

### Total RNA extraction

Total RNA was isolated from germinating yeast spore cultures by the “mechanical disruption protocol” using RNeasy MIDI kit (for RNA-Seq) or RNeasy mini kit (for Real-Time qRT-PCR) (Qiagen), following the manufacturer’s instructions. RNA integrity and concentration were assessed using the Bioanalyzer 2100 with the RNA 6000 Nano Lab Chip kit (Agilent) and the ND-1000 UV-visible light spectrophotometer (NanoDrop Technologies).

### RNA-Seq

Poly-A mRNA was enriched from ∼10 µg of total RNA with oligo dT beads (Invitrogen). ∼100 ng of poly-A mRNA recovered was used to construct multiplexed strand-specific RNA-Seq libraries as per manufacturer’s instruction (NEXTflexTM Rapid Directional RNA-SEQ Kit, dUTP-Based, v2). Individual library quality was assessed with an Agilent 2100 Bioanalyzer and quantified with a QuBit 2.0 fluorometer before pooling for sequencing on a HiSeq 2000 (1x101 bp read) yielding an average read count of 14,009,863 per sample with an average overall alignment: 95.23%. The pooled libraries were quantified using the KAPA quantification kit (KAPA Biosystems) prior to cluster formation. The accession number for the RNA-Seq data reported in this paper is NCBI: GSE110326.

### Differential Expression Analysis

The raw Fastq reads were trimmed for adapter sequences and low-quality bases using Trimmomatic (v.0.33) (*36*) (parameters: LEADING:3 TRAILING:3 SLIDINGWINDOW:4:15 MINLEN:36). These trimmed raw reads were then aligned to the Yeast genome (*Saccharomyces cerevisiae* strain SK1) using hisat2(*37*) (v.2.0.4) (parameters: --rna-strandness R) and the corresponding Ensemble annotation file (Saccharomyces_cerevisiae R64-1-1.86 gtf-file). The resulting sam files were then converted to bam files that were used to generate feature read counts using Python package based htseq-count of HTSeq (*38*) (v.0.6.1p1) (parameters: default union-counting mode, --stranded=reverse). The HTSeq read counts were used for the differential expression (DE) analysis using the edgeR (*39*) package (available in R (v.3.1.3)). A false discovery rate (FDR) cutoff of 0.05 and fold-changes were used to sift the genes that were significantly differentially expressed (DE). The GO terms of the DE genes were analyzed by DAVID (*40*) and visualized using REVIGO (*27*).

### Real-Time qRT-PCR

One microgram of total RNA was reverse-transcribed into cDNA using QuantiTect Reverse Transcription Kit (Qiagen). As a control for genomic DNA contamination, the same reaction was performed without Reverse Transcriptase. Real-time PCR was performed in a final volume of 20 µl containing 20 ng of cDNA using SYBR Fast Universal qPCR Kit (Kapa Biosystems) and analyzed using the Quant Studio 6 Flex system (Applied Biosystems). The real-time PCR conditions is one hold at [95°C, 180 secs], followed by 40 cycles of [95°C, 1 sec] and [60°C, 20 sec] steps. After amplification, a melting-curve analysis was done to verify PCR specificity and the absence of primer dimers. For quantification, the abundance of each gene was determined relative to the house keeping transcript *ACT1* and the relative gene expression between the DMSO control and Rapamycin-treated conditions were calculated using the 2^-ΔΔCt^ method (*41*). A list of primers used for Real-Time qRT-PCR is provided in Supplementary Table S4.

### Phospho-PKA substrate Western blot analysis

For phospho-PKA substrate western blots, protein extracts were prepared from trichloroacetic acid -treated cells. Exponentially growing cells were treated with DMSO or Rapamycin. Then, the cells were collected and suspended in 250µl of 10% trichloroacetic acid. Next, the cells were disrupted with glass beads on Precellys ® 24 homogenizer (Bertin Technologies). After that, the trichloroacetic acid-pellets were resuspended in 100 µl of 2× SDS gel-loading buffer plus 50 µl of 1M Tris base to adjust the pH. Samples were then boiled for 10 min and followed by centrifugation at 14000×g for 5 min. The supernatants were separated on 8% SDS-PAGE gels and transferred onto nitrocellulose membranes. Blots were then blocked with 5% milk in PBS/0.1% Tween 20. For phospho-PKA substrate, blots were probed with Phospho-PKA Substrate (RRXS*/T*) antibody (1:1000; cat no. 9624, cell signaling technology) followed by goat anti-rabbit antibody conjugated with horseradish peroxidase (1:5000; Santa-Cruz). For β-actin, blots were probed with anti-beta actin antibody (1:5000; cat no. ab8224, abcam) followed by goat anti-mouse antibody conjugated with horseradish peroxidase (1:5000; Santa-Cruz). Finally, the blots were developed with ECL prime western blotting detection reagent (Amersham Pharmacia Biotech).

### Sporulation

Yeast strains were streak purified on YPD plates (1% yeast extract, 2% peptone, 2% glucose and 2% agar). At least five colonies were patched onto YPD plates and incubated at 30 °C for 24 h. Cells were then patched onto sporulation plates (0.82% sodium acetate, 0.19% potassium chloride, 0.035% magnesium sulphate, 0.12% sodium chloride and 1.5% agar) and YPD plates and incubated at 30 °C for 24 h. Sporulation efficiency was examined using a light microscope and the corresponding YPD patch from the colony that sporulated best was inoculated in 50 ml YPD liquid media (250 ml conical flask). Cells were grown in a 30 °C shaker at 220 rpm for 16 h. Cells were then washed twice with sporulation medium (0.25% yeast extract, 1.5% potassium acetate, 0.25% glucose). Cells (3 OD_600_/ml) were resuspended in 50 ml of liquid sporulation media (500 ml conical flask). Sporulation was induced for three and half days in the 30 °C shaker at 220 rpm.

### Spore purification

Spore purification was carried out as previously described with slight modifications (*25*). Briefly, the sporulation culture was centrifuged at 1000 g for 5 min. Cells (40 OD600/ml) were resuspended in softening buffer (10 mM dithiothreitol, 100 mM Tris-SO4, pH 9.4) and incubated for 15 min with shaking at 30 °C. Cells were pelleted and resuspended in spheroplasting buffer (2.1 M sorbitol, 10 mM potassium phosphate, pH 7.2). Zymolyase-20T (cat no. 320921, MP Biomedicals, LLC) was added to the cells at a concentration of 2 mg/ml. Spheroplasting reaction was performed for 30 min with shaking at 30 °C. The suspension was washed thrice and resuspended in spheroplasting buffer. The spore suspension was sonicated (Amplitude 40 %, Cycle 10 of 20 sec with 10 sec interval) briefly to disperse the spores and kept on ice. Fresh spores were used for all the experiments.

### Spore germination

Purified spores were suspended at approximately 1 OD600 cells/ml YPD medium or 2% glucose and incubated at 30 °C with 220 rpm shaking. Samples were taken for analysis at different time points including the zero-time point. To examine the germination under different conditions purified spores were incubated with chemicals in YPD medium or 2% glucose and sampling was performed over different time points.

### Microscopy

Spore cultures at different time points were fixed by 4% paraformaldehyde for 15 minutes with shaking at room temperature. Fixed spores were washed and resuspended in 100 mM K-phosphate buffer (pH 7.5) / 2 M sorbitol. Spores were briefly sonicated and directly visualized under either bright-field or fluorescence microscope.

## SUPPLEMENTARY MATERIALS

Table S1. Comparative transcriptomic analysis of purified spores and spores transferred to nutrient medium.

Table S2. Comparative transcriptomic analyses of spores transferred to nutrient medium and spores transferred to nutrient medium containing rapamycin (2 µM).

Table S3. A list of yeast strains used in the study.

Table S4. A list of primers used for Real-Time qRT-PCR analyses in the study.

Table S5 Comparative transcriptomic analysis of purified spores and spores transferred to nutrient medium containing rapamycin (2 µM)

Table S6. Lists of genes in the 10 categories reported in a transcriptomic study of spore germination in yeast (*1*).

## Acknowledgements

We thank Prof. James Broach (Princeton University, USA), Prof. Claudio de Virgilio (University of Fribourg, Switzerland), Prof. Mike Hall (Biozentrum University of Basel, Switzerland), Prof. Paul Herman (The Ohio State University) and Prof. Kim Nasmyth (University of Oxford) for sharing plasmids and yeast strains with us. We would also like to thank Prof. Uttam Surana (IMCB, Singapore) and Dr Dmitry Ivanov (IBS, South Korea) for comments on the manuscript.

## Author contributions

MA performed the spore germination assays, TORC1 activation experiments and Gene Ontology analysis of RNA-Seq data. JHW helped in preparing the RNA for RNA-Seq experiments and performed the qRT-PCR experiments. YCL performed TORC1 activation assay. KWK constructed the libraries for NGS analysis. QFC constructed yeast strains and CJHG helped with the RT-PCR experiments. VGK did the bioinformatic analysis of RNA-Seq data. SH supervised all aspects of RNA-Seq analysis. PA conceived, supervised the overall project and wrote the manuscript. All authors commented on the manuscript and agreed to its final version.

## Data and materials availability

RNA-Seq data reported in this paper are publicly available at NCBI (Accession number GSE110326). All strains used in the study are available upon request.

## Legends for Supplementary figures

**Supplementary Figure S1.**
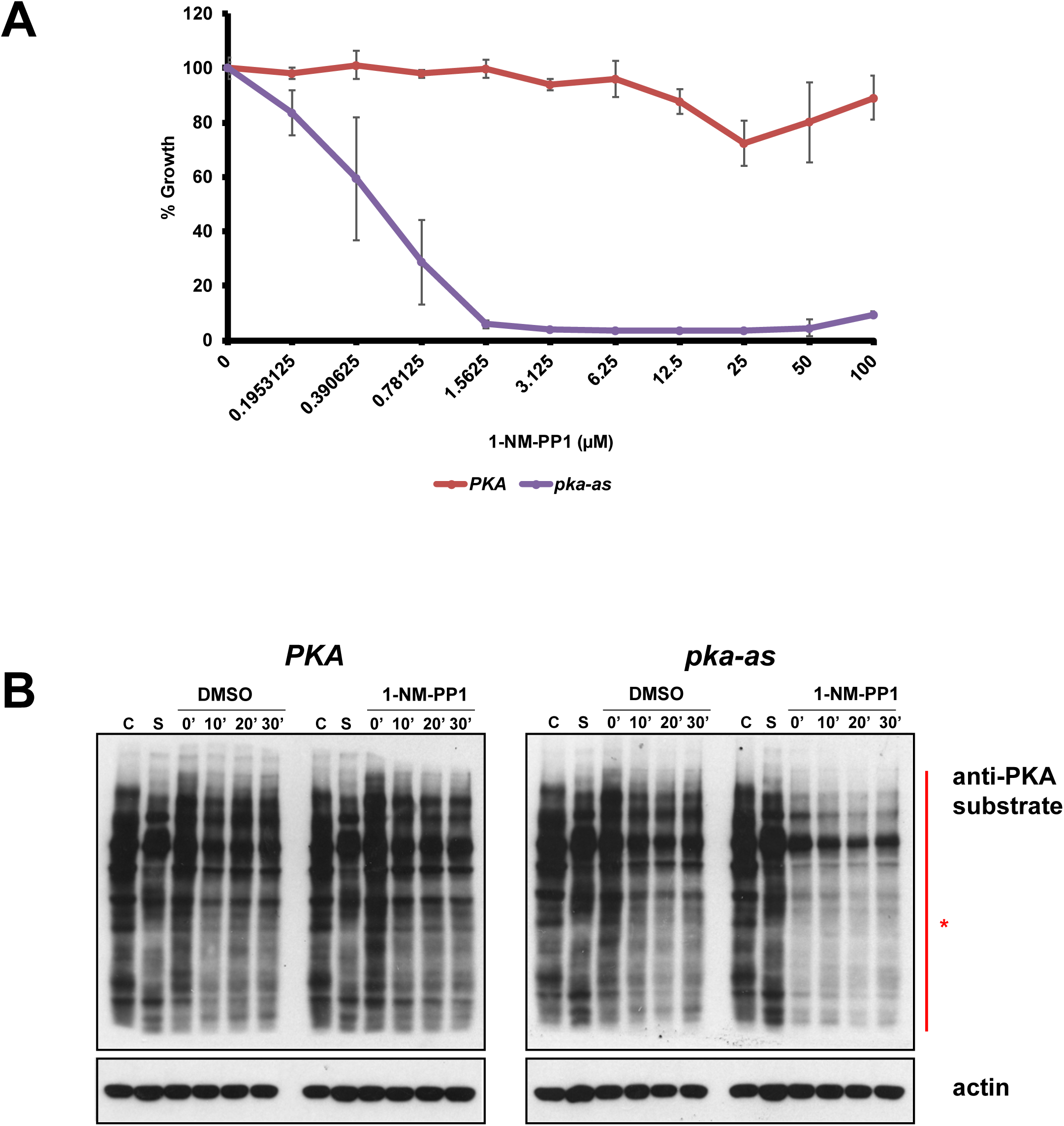
1-NM-PP1 inhibits the growth and PKA activity in *pka-as* but not in wild type cells. A) Wild type (PKA) and *pka-as* yeast cultures were diluted to a starting OD=O.2 in YPD medium containing either DMSO or 1-NM-PP1 at the various concentrations indicated and incubated at 30 °C in a shaker (250 rpm). Normalized growth after 24 hours of incubation at 30 °C is plotted for the various cultures. B) Wild type and *pka-as* cells subjected to complete nutrient starvation were transferred to 2% glucose solution in the presence of either DMSO or rapamycin (2 µM) or 1-NM-PP1 (25 µM). Aliquots of the cultures were taken after 0’, 10’, 20’ and 30’ and used for preparing protein extracts. Protein samples were analyzed by Western blotting using anti-PKA substrate and anti-actin antibodies.

**Supplementary Figure 2.**
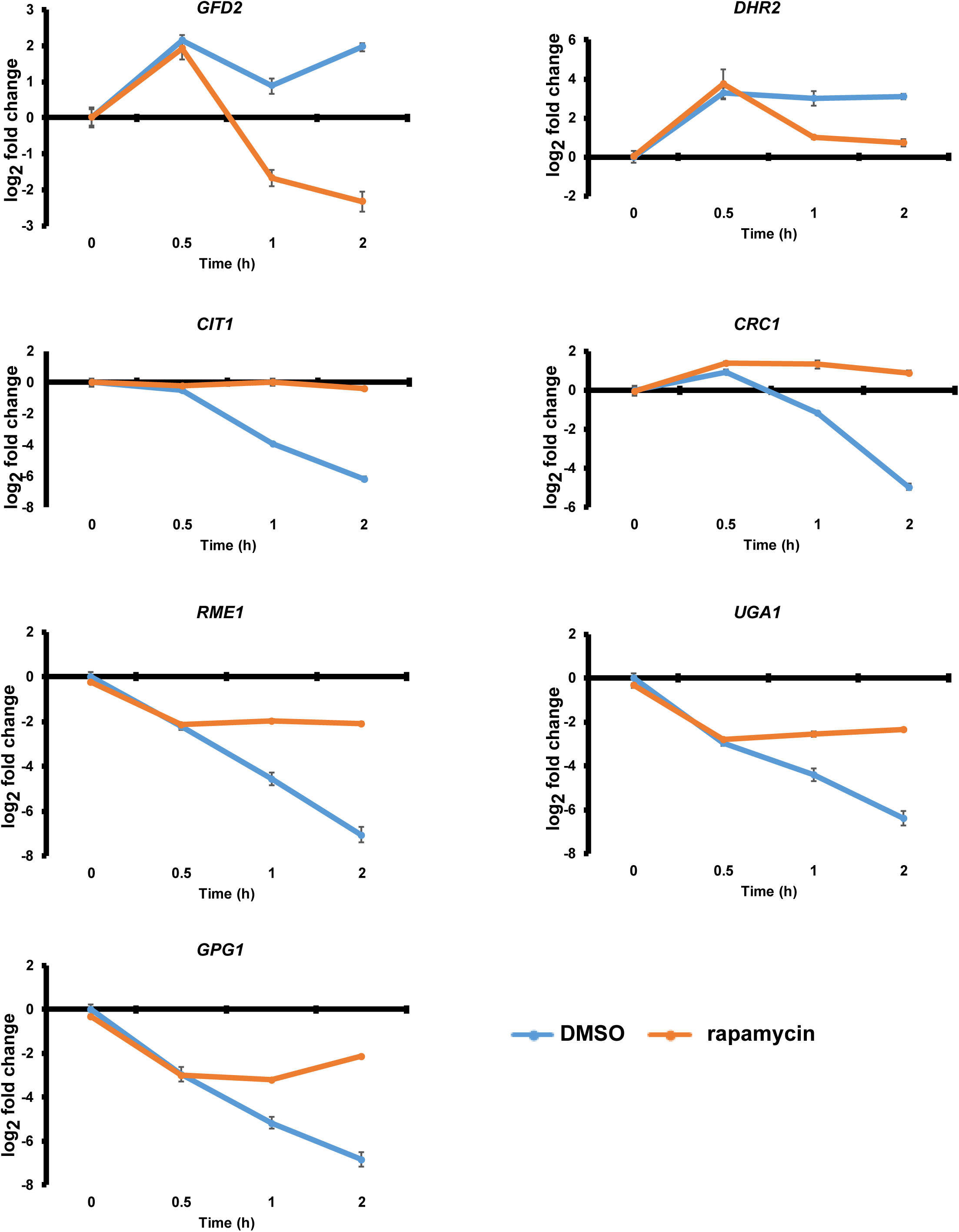
TORC1 is required to maintain the expression of TGC genes in glucose-containing growth medium. Wild type cells grown into mid-log phase in YPD medium were treated with either DMSO or rapamycin (200 nM). Aliquots of the cultures were taken after 0, 0.5, 1 and 2 hours. RNA was extracted from the cultures and the expression of the TGC genes was analyzed by Real-Time qRT-PCR.

**Supplementary Figure S3.**
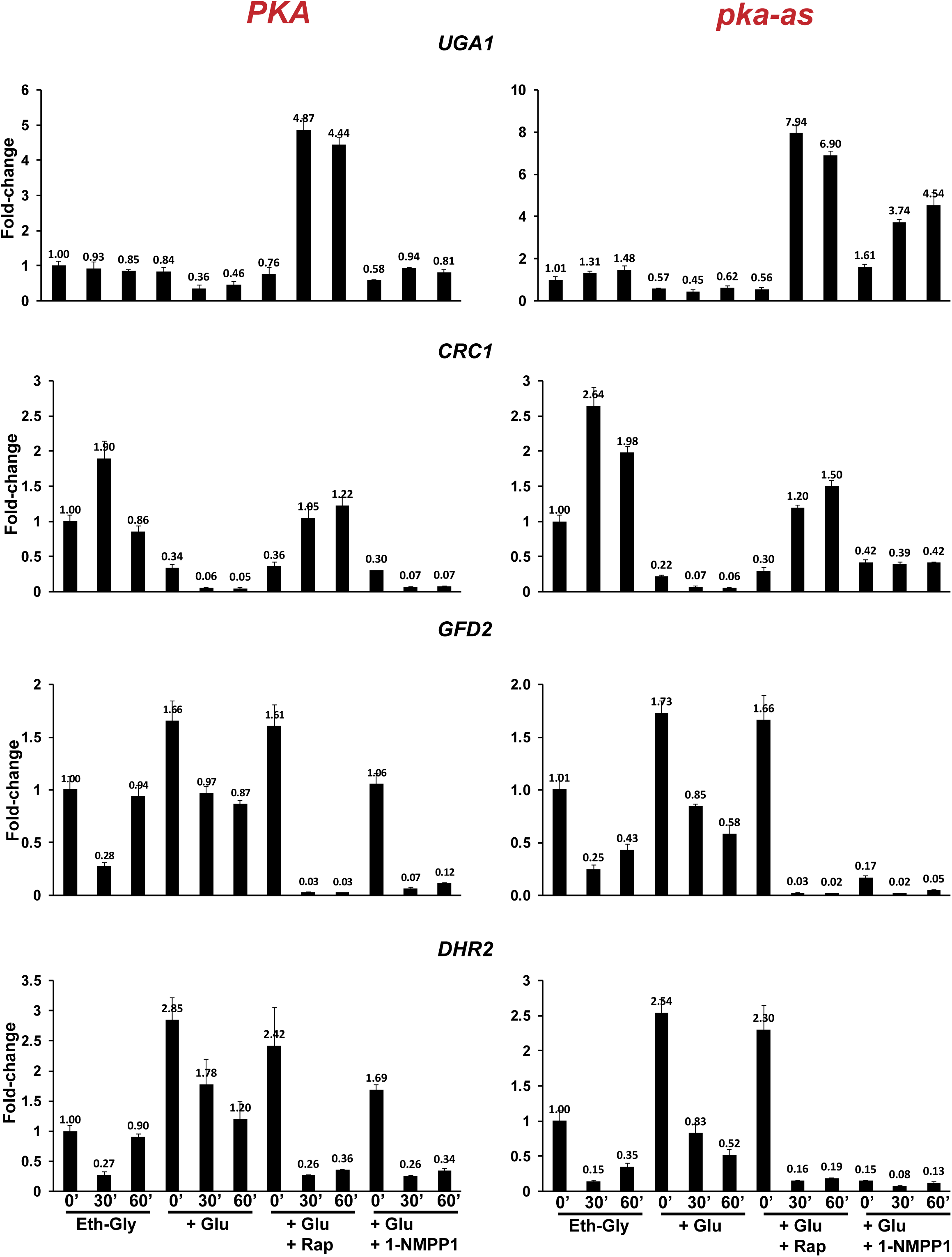
TORC1 and PKA co-regulate the expression of glucose-responsive genes. Wild type (PKA) and *pka-as* cells were grown to logarithmic phase and then treated with either rapamycin (200 nM) or 1-NM-PP1 (20 µM) or DMSO. Aliquots of the cultures were taken after 0, 30’ and 60’. RNA was extracted from the cultures and the expression of the 4 glucose response genes (*GFD2*, *UGA1*, *CRC1* and *DHR2*) were analyzed by Real-Time qRT-PCR.

**Supplementary Figure S4.**
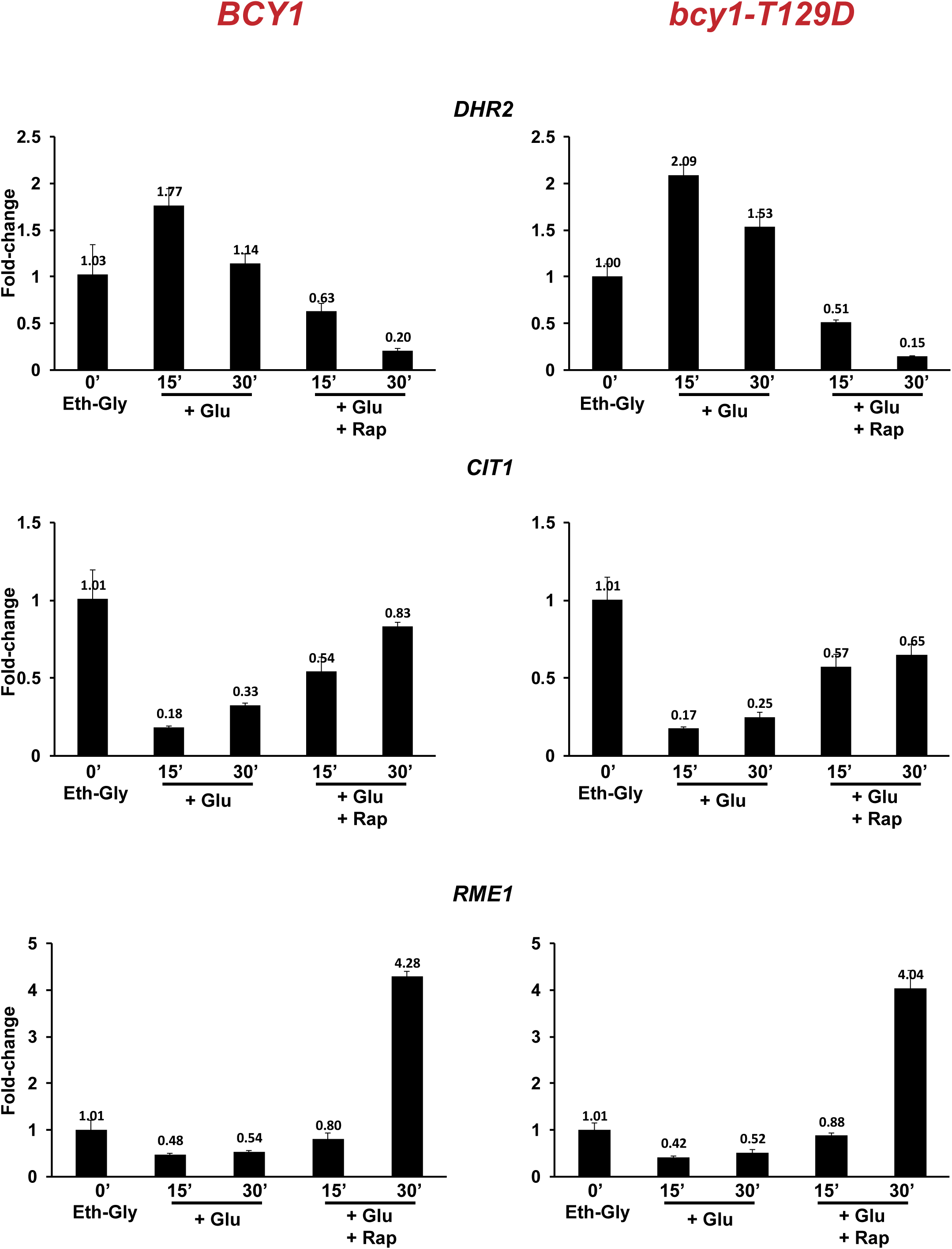
TORC1 regulates the glucose-responsive genes independently of Bcy1 T129 dephosphorylation. Cells expressing wild type Bcy1 and mutant *bcy1-T129D* were grown to logarithmic phase in SC-EG medium. Glucose was added to the cultures at the final concentration of 2% along with either rapamycin (200 nM) or DMSO. Aliquots of the cultures were taken after 0, 15’ and 30’. RNA was extracted from the cultures and the expression of the glucose response genes *DHR2*, *CIT1* and *RME1* were analyzed by Real-Time qRT-PCR.

**Supplementary Figure S5.**
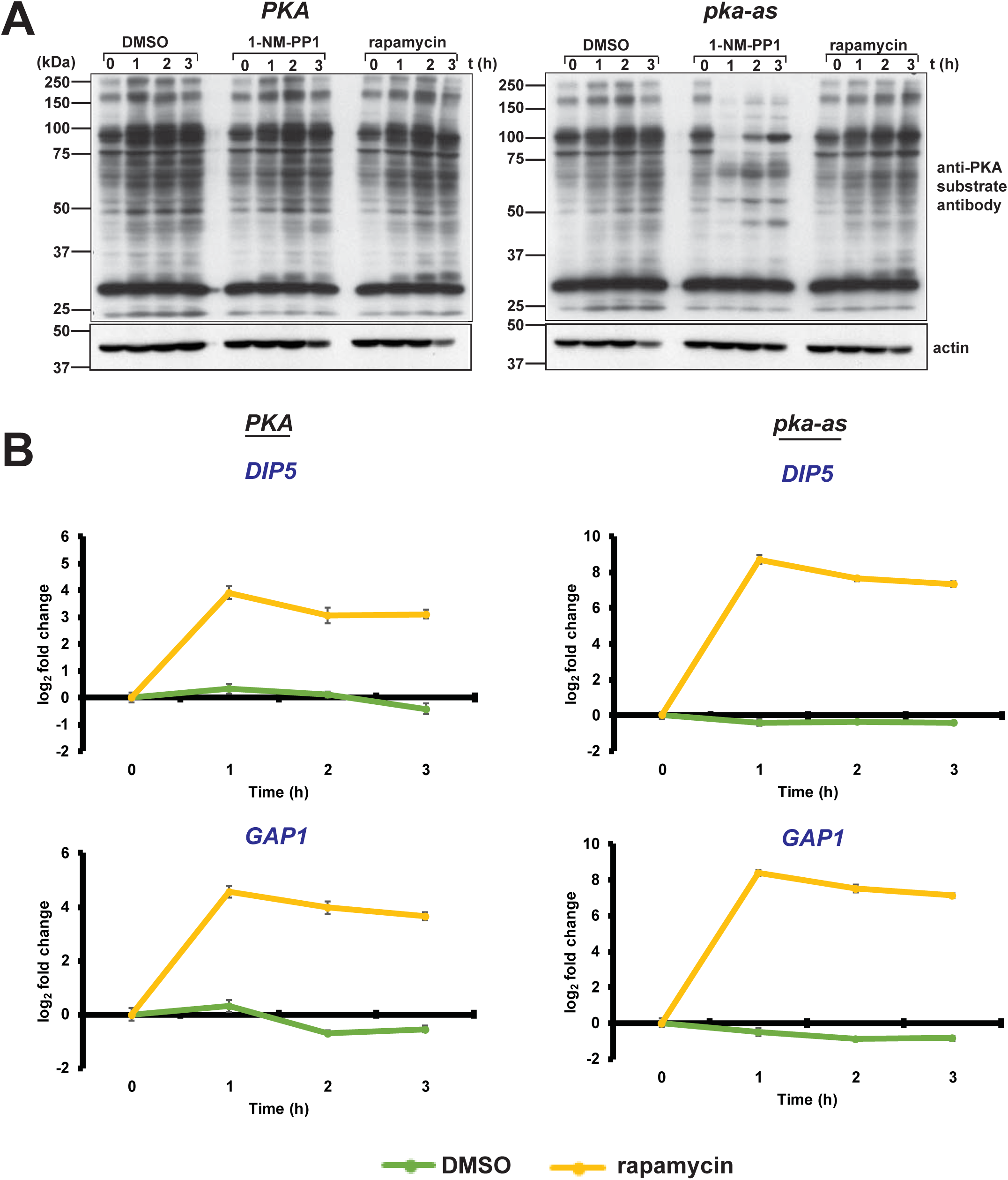
Inhibition of TORC1 has no detectable effect on overall PKA activity. A) Wild type cells (PKA) and *pka-as* cells (containing an analogue-sensitive allele of PKA) were grown into mid-log phase and were then treated with either DMSO or 1-NM-PP1 (25 µM) or rapamycin (200 nM). Whole cell extracts were prepared from aliquots of cells were taken after 0, 1, 2 and 3 hours and were analyzed by Western blotting using an anti-PKA substrate antibody and actin antibody (loading control). B) Expression of TORC1 target genes *DIP5* and *GAP1* was assayed for the experiment in A by Real-Time qRT-PCR analysis. Levels of transcripts were normalized with respect to actin mRNA.

**Supplementary Figure S6.**
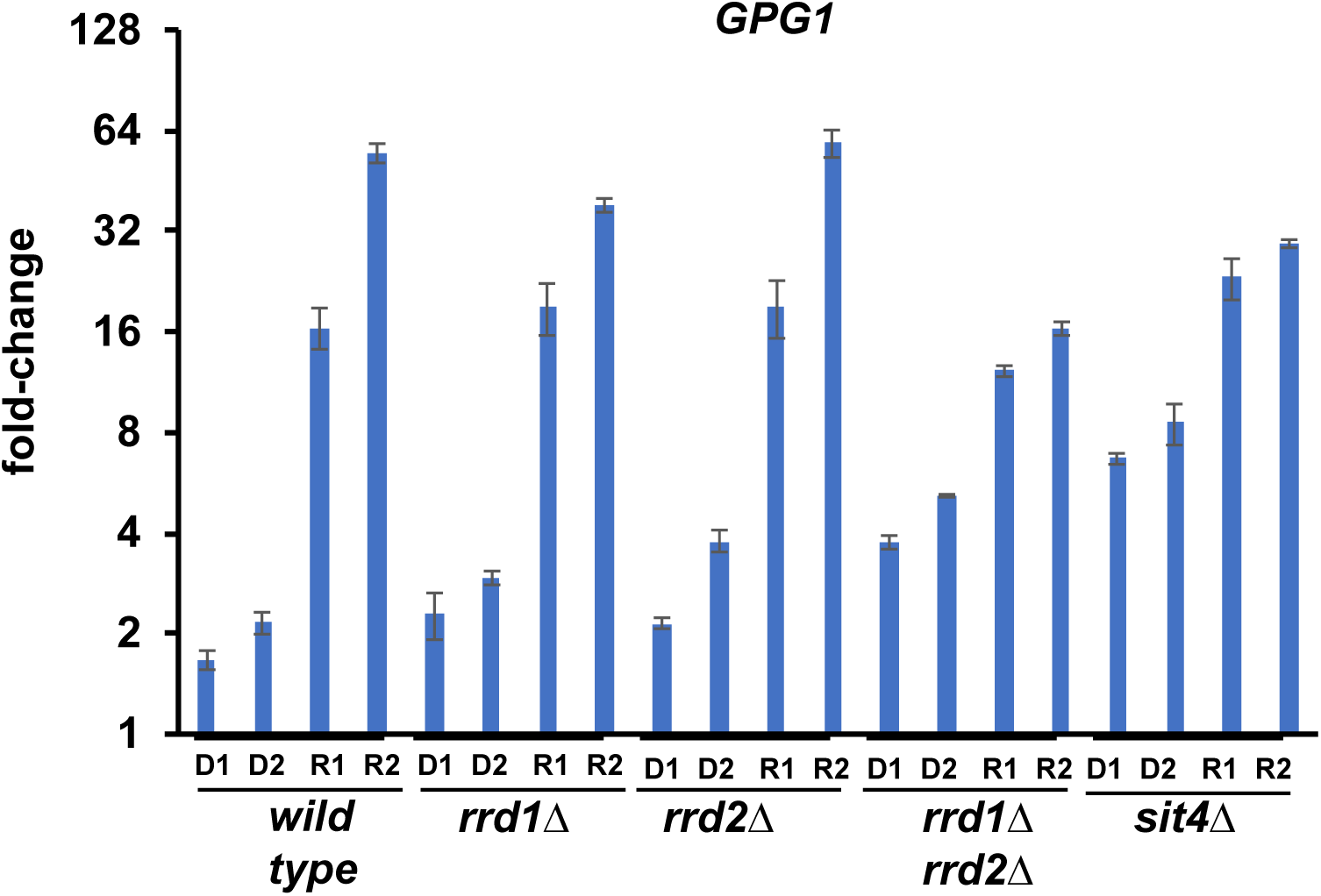
TORC1 regulates the glucose-responsive genes by inhibiting Sit4 and Rrd1/Rrd2 proteins. Wild type, *sit4Δ, rrd1Δ, rrd2Δ* and *rrd1Δ rrd2Δ* cells were grown to logarithmic phase and then treated with either rapamycin (200 nM) or DMSO. Aliquots of the cultures were taken after 0’, 30’ and 60’. RNA was extracted from the cultures and the expression of the 7 glucose response genes (*GFD2*, *GPG1*, *UGA1*, *RME1*, *CIT1*, *CRC1* and *DHR2*) were analyzed by Real-Time qRT-PCR. Results for *GPG1* are presented here. D1/R1 and D2/R2 represent samples from DMSO-treated/ Rapamycin-treated cells after 30’ and 60’ respectively and the expression fold-change values were normalized with respect to DMSO-treated cells at t=0’.

**Supplementary Figure S7.**
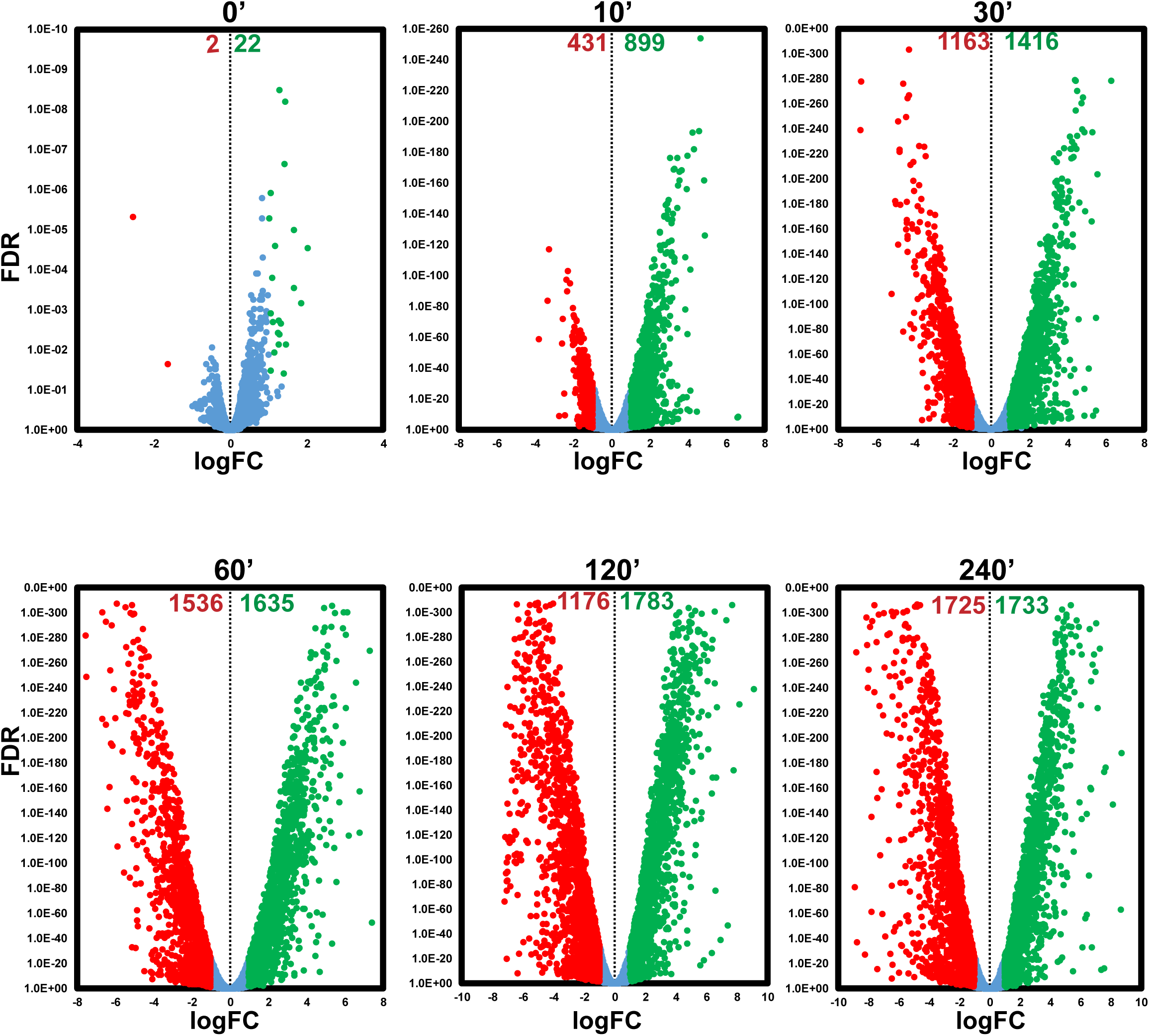
Transcriptomic analysis of spore germination. For each time point the RNA seq data of germinating spore cultures were compared with that of purified spores and the logFC (Fold Change) was plotted against False Discovery Rate (FDR). Differentially Expressed Genes (at least 2-fold difference in comparison to the spore) were identified. Positively and negatively regulated genes during spore germination are indicated by green and red dots respectively and their numbers are indicated at the top of the plot. Unaffected genes are represented by blue dots.

**Supplementary Figure S8.**
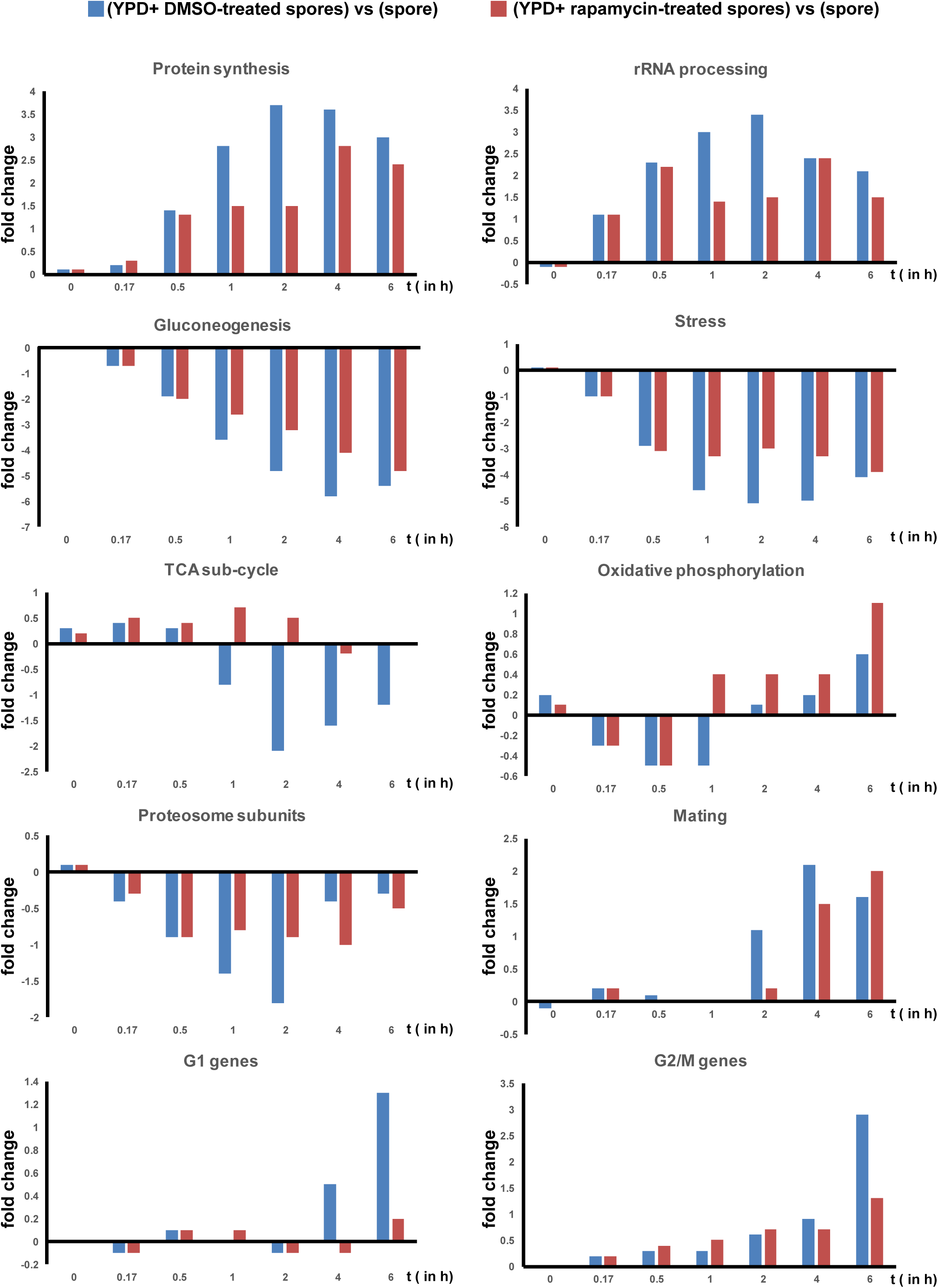
Comparative gene expression analysis of specific gene modules during spore germination in the presence and absence of rapamycin. Expression of genes in 10 specific modules described in an earlier transcriptomic study of spore germination (Joseph-Strauss et al Genome Biology2007 8: R241) was examined in our RNA-SEQ data. The transcripts levels of genes in 10 modules in spores incubated in either YPD + DMSO or YPD + rapamycin, for 0, 0.17, 0.5, 1, 2, 4 and 6 h was compared with the corresponding level in ungerminated spores. Blue and red bars indicate the fold-change values for ‘YPD + DMSO’ and ‘YPD + rapamycin’ cultures respectively. Comparison of gene expression between ‘spores’ and ‘spores + YPD + rapamycin’ are shown in Supplementary Table S5. Lists of genes in the 10 categories are available in Supplementary Table S6.

**Supplementary Figure S9.**
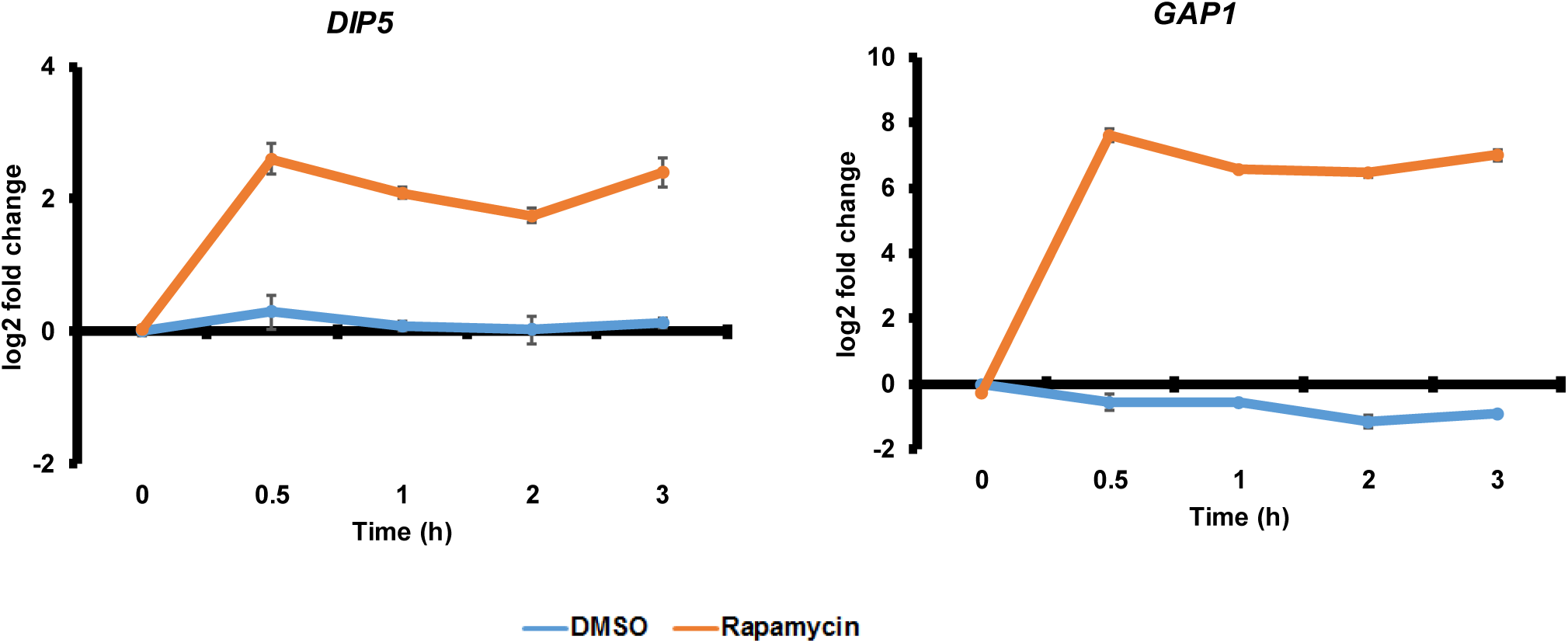
Validation of RNA-Seq data by Real-Time qRT-PCR analysis of TORC1 target genes *DIP5* and *GAP1*. Spores were transferred to YPD medium with either DMSO or rapamycin (2 µM). Aliquots of yeast cells taken at the indicated time points (0, 0.5, 1, 2 and 3 h) from the two cultures were used for preparing RNA. Expression of TORC1 target genes (*DIP5* and *GAP1*) was assayed by Real-Time qRT-PCR analysis. Levels of transcripts were normalized with respect to actin mRNA.

**Supplementary Figure S10.**
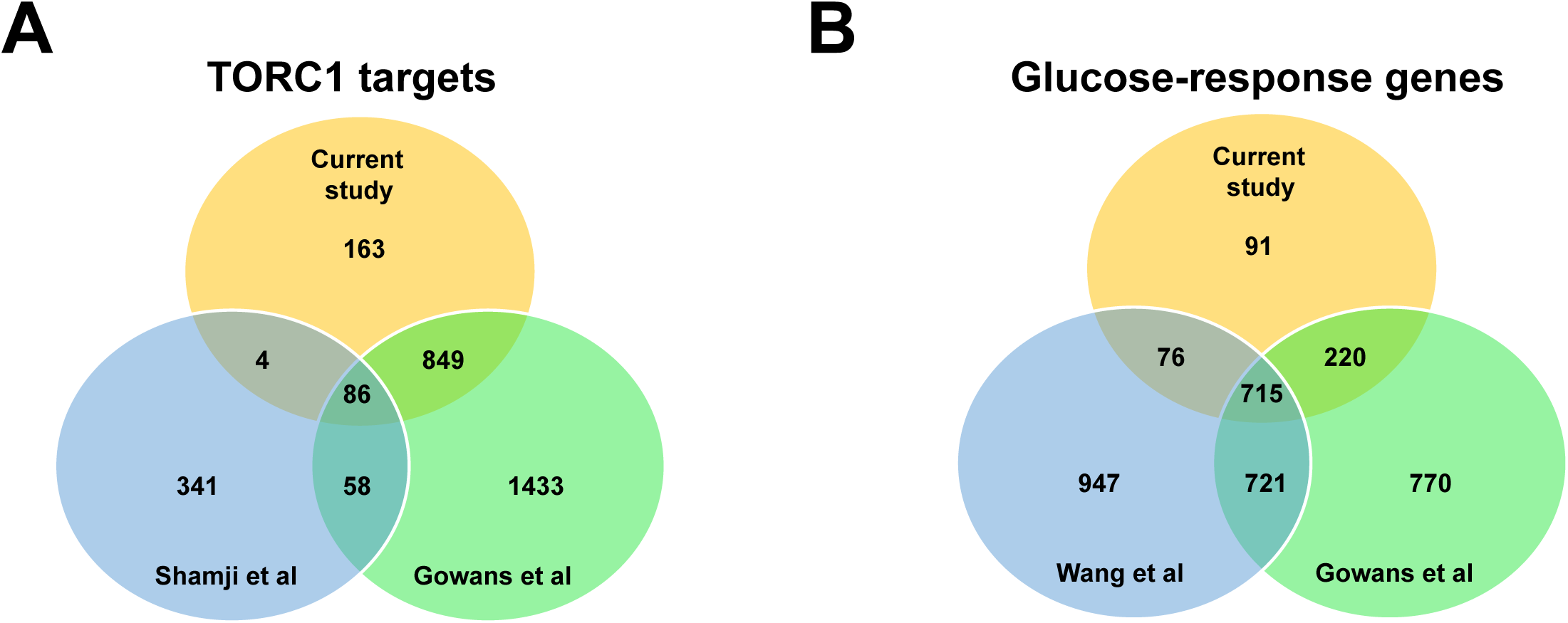
Overlap of Glucose-responsive genes with TORC1 target genes. A) Comparison of our TORC1 target list identified during spore germination with those reported in a Micro-array-based study (Shamji et al. (*28*)) and an RNA-Seq study (Gowans et al. (*19*)) both performed with vegetative cells. B) Comparison of the list of glucose-responsive genes (Wang et al. (*14*)) with TORC1 target lists identified by our RNA-Seq analysis and an independent RNA-Seq study (Gowans et al.) (*19*).\

**Supplementary Figure S11.**
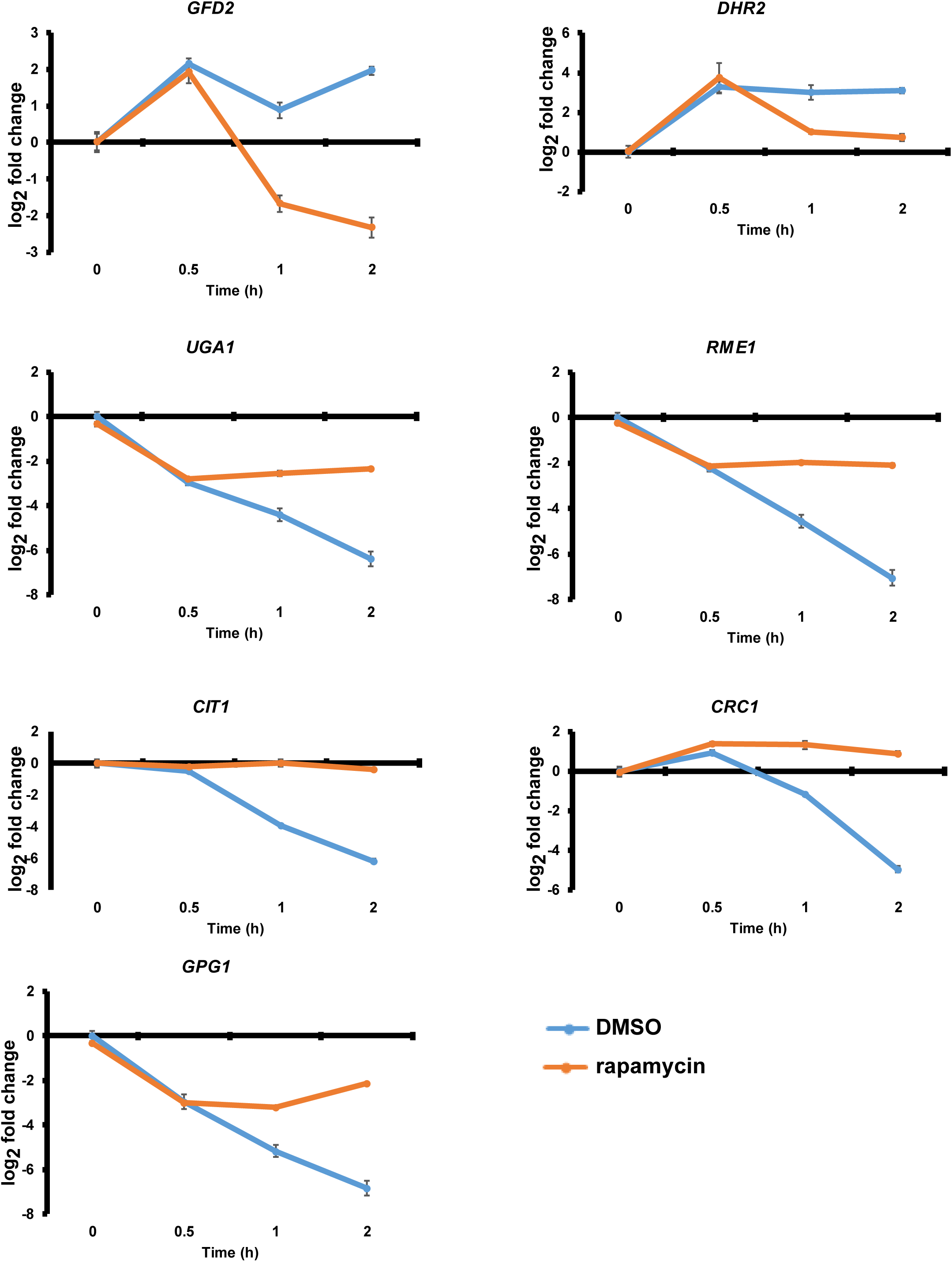
TORC1 regulates a subset of glucose-responsive genes during spore germination. Spores were transferred to YPD medium with either DMSO or rapamycin (2 µM). Aliquots of yeast cells taken at the indicated time points (0 h, 0.5 h, 1 h, and 2 h) from the two cultures were used for preparing RNA. Expression of 7 glucose-responsive genes (*GFD2*, *GPG1*, *UGA1*, *RME1*, *CIT1*, *CRC1* and *DHR2*) was assayed by Real-Time qRT-PCR analysis. Levels of transcripts were normalized with respect to actin mRNA.

**Supplementary Figure S12.**
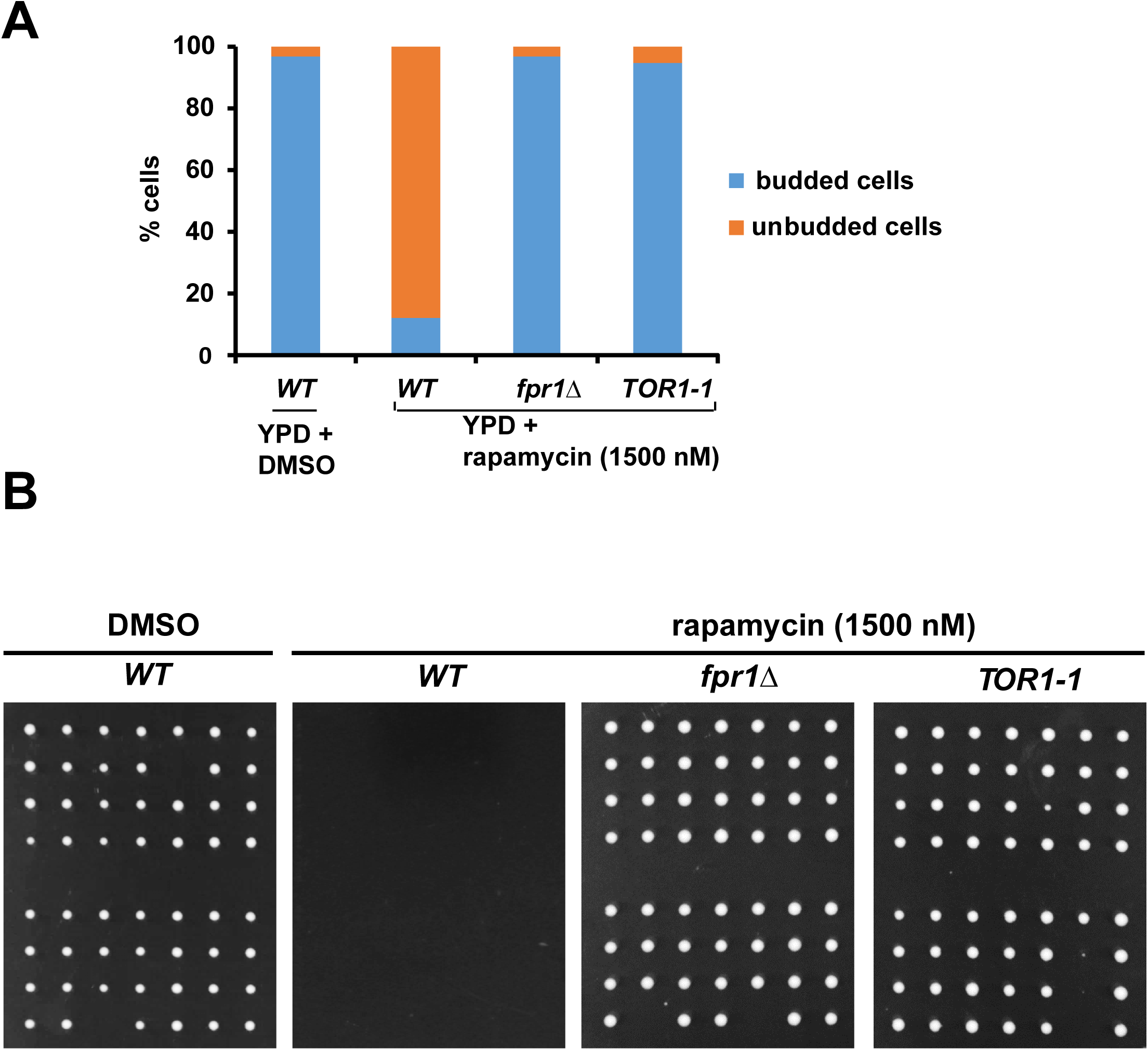
TORC1 is required for spore germination. A) Fourteen asci resulting from sporulation of wild type or *fpr1Δ* or *TOR1-1* diploid cells were dissected on YPD + agar plates containing 1.5 µM rapamycin. Fourteen asci from wild type diploid cells were also dissected on a YPD agar plate without rapamycin. Percentage of budded cells was calculated by examining the spore morphology under the dissection microscope after 6 hours following dissection and is indicated in the plot. B) Images of the agar plates described in A following incubation at 30 °C for 2 days are presented.

**Supplementary Figure S13.**
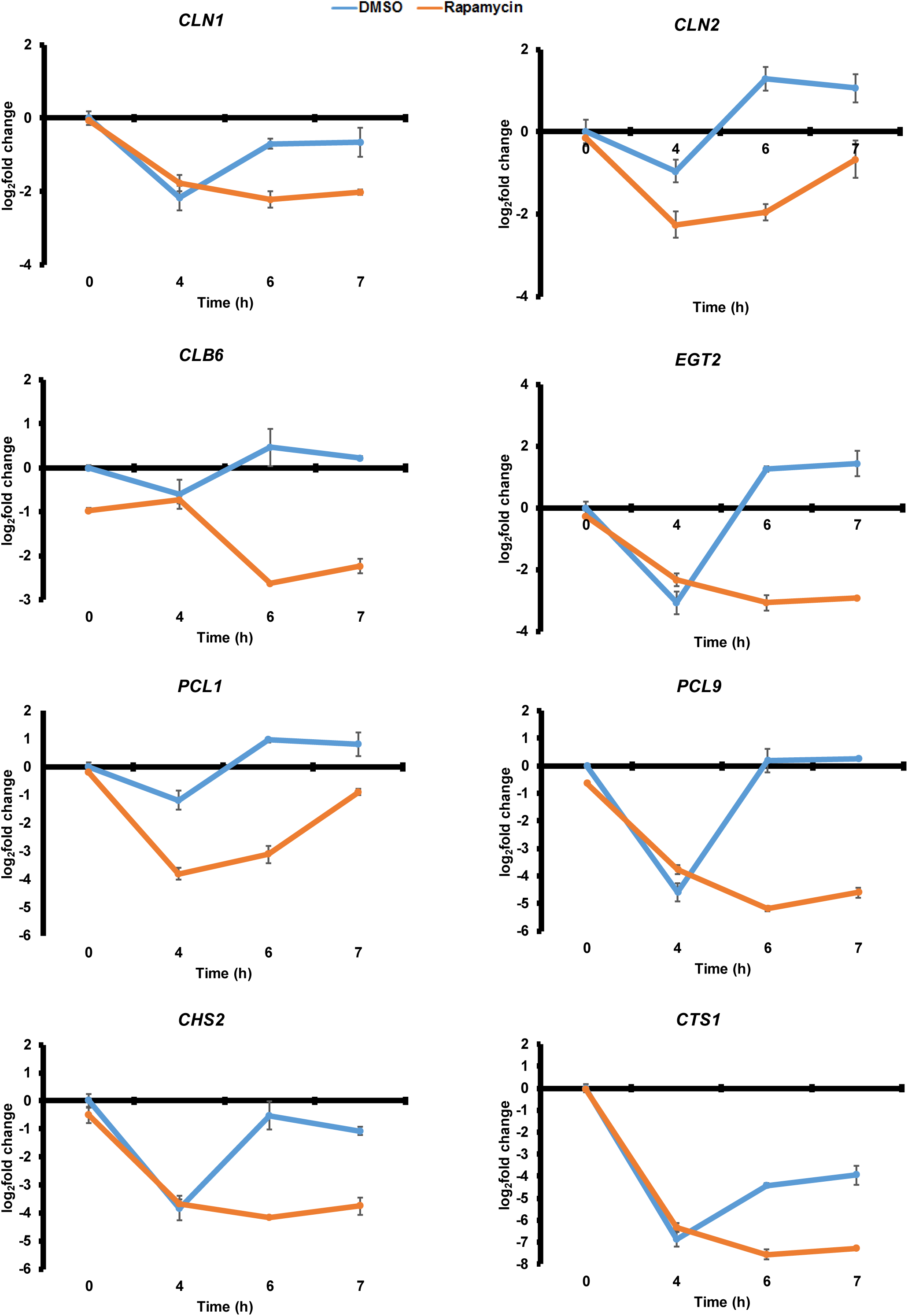
TORC1 is required for the expression of cell cycle genes during spore germination. Spores were transferred to YPD medium with either DMSO or rapamycin (2 µM). Aliquots of yeast cells taken at the indicated time points (0, 4, 6 and 7 h) from the two cultures were used for preparing RNA. Expression of cell cycle genes *CLN1*, *CLN2*, *CLB6*, *EGT2*, *PCL1*, *PCL9*, *CHS2* and *CTS1* was assayed by Real-Time qPCR analysis. Levels of transcripts were normalized with respect to actin mRNA.

**Supplementary Figure S14.**
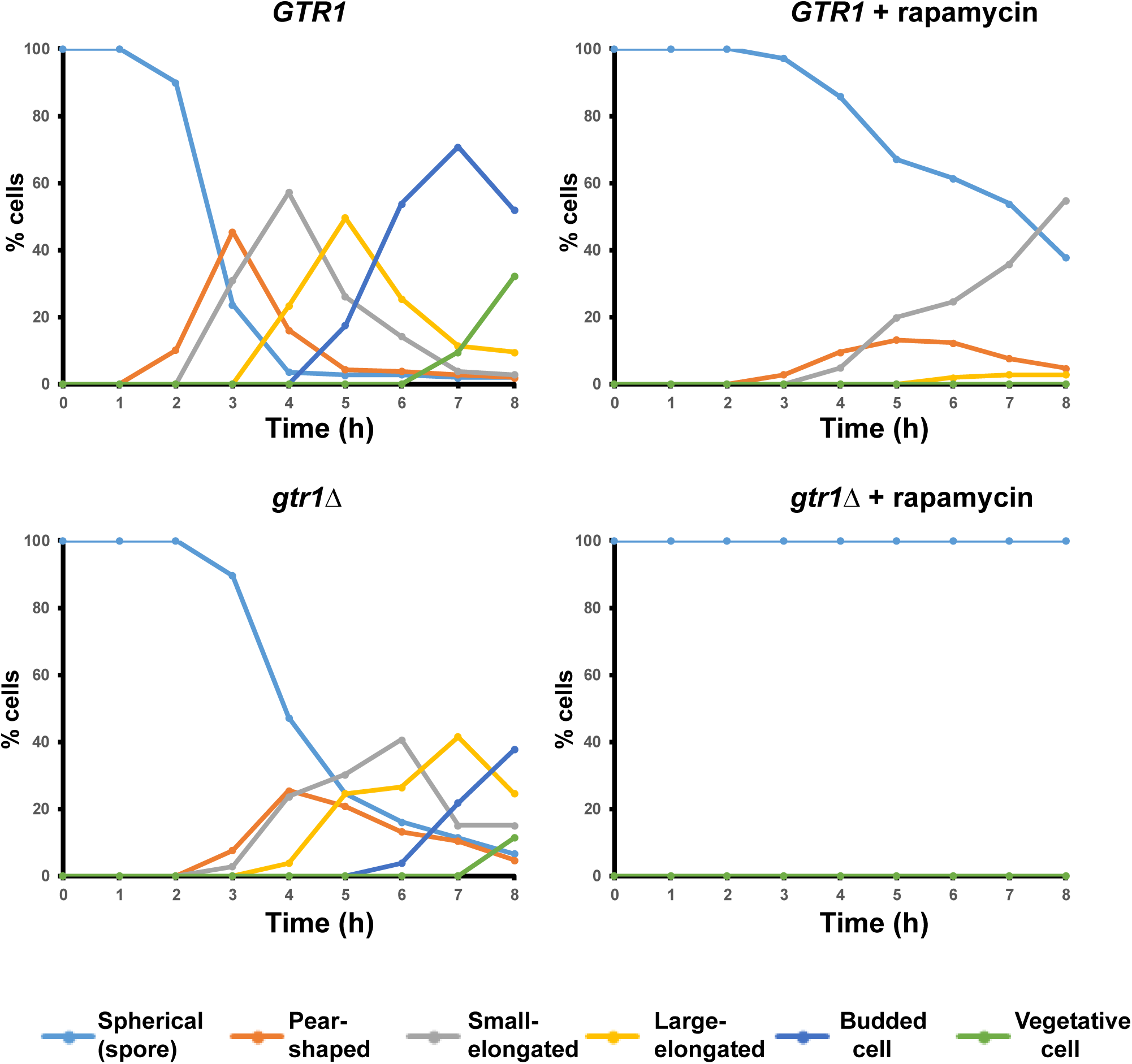
*gtr1Δ* spores fail to germinate in the presence of rapamycin. Wild type and *gtr1Δ* spores were transferred into nutrient medium in the presence of either DMSO or rapamycin (2 µM). Progress of spore germination was assayed by scoring fraction of cells with different morphologies at the indicated time points for 8 hours following transfer into YPD medium.

**Supplementary Figure S15.**
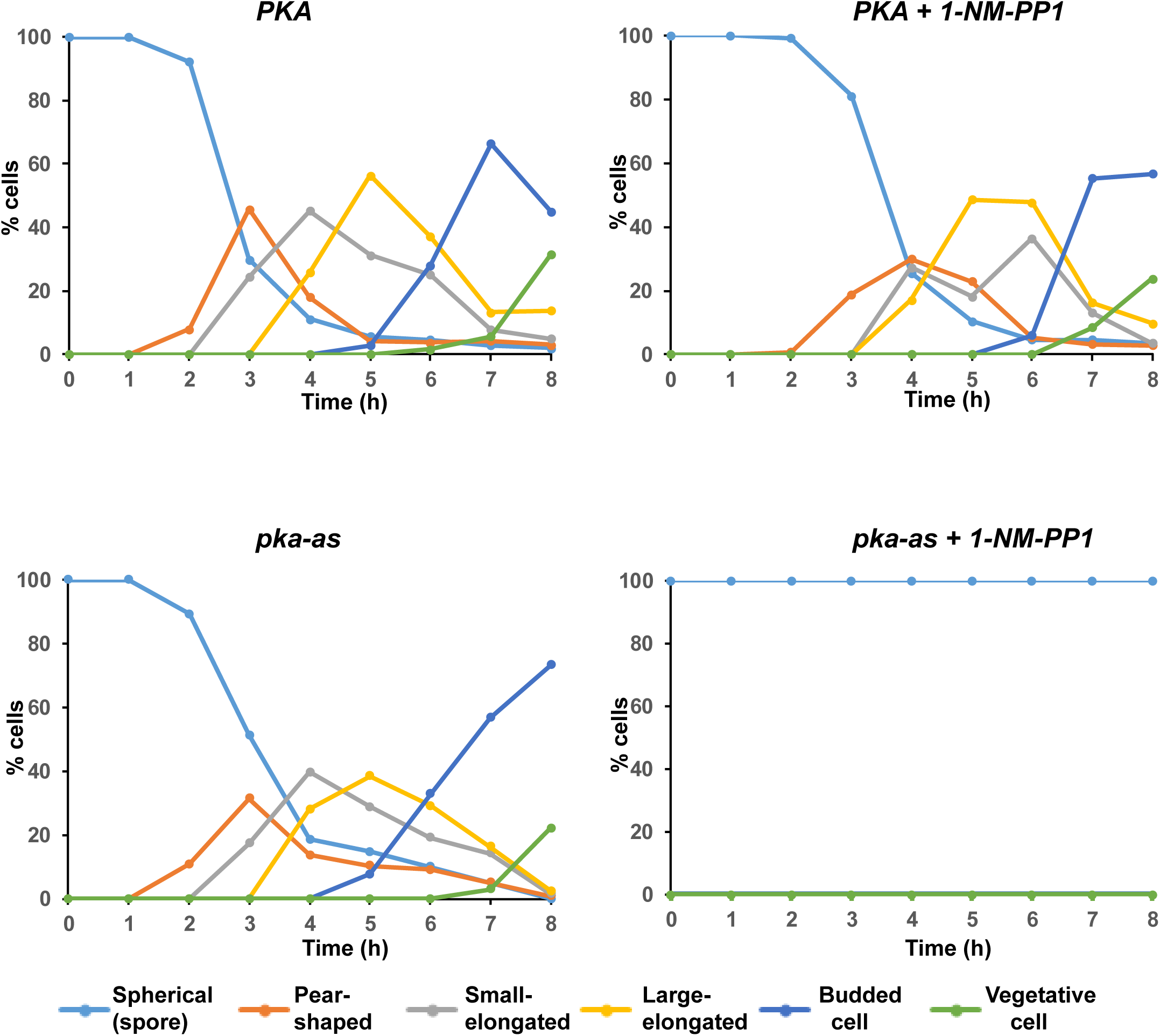
PKA is required for spore germination. Wild type and *pka-as* spores were transferred into nutrient medium in the presence of either DMSO or 1-NM-PP1 (25 µM). Progress of spore germination was assayed by scoring fraction of cells with different morphologies at the indicated time points for 8 hours following transfer into YPD medium.

