## Supplementary material for "TORC1 regulates the transcriptional response to glucose and developmental cycle via the Tap42-Sit4-Rrd1/2 pathway in *Saccharomyces cerevisiae*": Table S3

**Table S3 List of strains used in the study**

All strains are derivatives of SK1 and have the following markers ho::LYS2 ura3 leu2 trp1 his3 lys2 All markers are homozygous in diploid strains unless otherwise mentioned

| Strain number | Genotype | Used in Figure |
| --- | --- | --- |
| 3526 | MATa *sch9: SCH9-HA6::KANMX6* | 1, 2, 4 |
| 3681 | MATa *gtr1:KANMX6* *sch9: SCH9-HA6::KANMX6* | 1, 2 |
| 1725 | MATa | 3, 5 |
| 3575 | MATa *sch9: SCH9-HA6::KANMX6 tpk1-as tpk2-as tpk3:KanMX6* | 4 |
| 3502 | MATa *sch9:KANMX6* | 5 |
| 3569 | MATa *tap42:KANMX6 TAP42-CEN- LEU2* | 6 |
| 3571 | MATa *tap42:KANMX6 tap42-11-CEN- LEU2* | 6 |
| 4860 | MATa *rrd1:NATMX6* | 7 |
| 3530 | MATa *rrd2:KANMX6* | 7 |
| 4861 | MATa *rrd1:NATMX6 rrd2:NATMX6* | 7 |
|  | MATa *sit4:HIS3MX6* | 7 |
| 1738 | MAT a/MATα | 1, 2A, 4,5,6,7, 13 |
| 3478 | MAT a/MATα *TOR1-1* | 1, 2A |
| 4154 | MATa/MATα *kog1: KOG1-3X-GFP: TRP1 vac7: VAC7-mCherry: NATMX6 sch9:SCH9-HA6::KANMX6* | 2B |
| 3555 | MATa/MATα *sch9: SCH9-HA6::KANMX6* | 2C, 3A, 3B, 12A |
| 3791 | MATa/MATα *gtr1:KANMX6 sch9: SCH9-HA6::KANMX6* | 3A, 3B, 12A |
| 3702 | MATa/MATα *gtr2:KANMX6 sch9: SCH9-HA6::KANMX6* | 3A |
| 3703 | MATa/MATα *ego1:KANMX6 sch9: SCH9-HA6::KANMX6* | 3A |
| 3526 | MATa *sch9: SCH9-HA6::KANMX6* | 1, 2 |
| 3576 | MATa *sch9: SCH9-HA6::KANMX6 tpk1-as tpk2-as tpk3:KanMX6* | 8, 12C |
| 4344 | MATa *bcy1:KANMX6 YCplac33::HA-BCY1 (CEN URA3)* | 9 |
| 4346 | MATa *bcy1:KANMX6 YCplac33::HA-bcy1--T129A (CEN URA3)* | 9 |
| 4347 | MATa *bcy1:KANMX6 YCplac33::HA- bcy1--T129D (CEN URA3)* | 9 |
| 4349 | MATa *bcy1: KANMX6YCplac33::HA-- bcy1--S145A (CEN URA3)* | 9 |
| 4351 | MATa *bcy1: KANMX6 YCplac33::HA-- bcy1--S145D (CEN URA3)* | 9 |
| 1725 | MATa | 10 |
| 3569 | MATa *tap42:KANMX6 TAP42-CEN- LEU2* | 11 |
| 3571 | MATa *tap42:KANMX6 tap42-11-CEN- LEU2* | 11 |
| 3681 | MATa *gtr1: KANMX6* *sch9: SCH9-HA6::KANMX6* | 12B |
| 3575 | MATa/MATα *sch9: SCH9-HA6::KANMX6 tpk1-as tpk2-as tpk3:KanMX6* | 13 |
