## Supplementary material for "TORC1 regulates the transcriptional response to glucose and developmental cycle via the Tap42-Sit4-Rrd1/2 pathway in *Saccharomyces cerevisiae*": Table S4

**Table S4. List of primers used for RT-qPCR**

| **Primer** | **Gene (S=forward & A=reverse)** | **Sequence (5' -> 3')** |
| --- | --- | --- |
| MA71 | ACT1_S | GAAGTGTGATGTCGATGTCC |
| MA72 | ACT1_A | TCTTTCTGGAGGAGCAATG |
| MA73 | GAP1_S | ATCCTTCCCACTTGTTATGG |
| MA74 | GAP1_A | GCCTTTTCTTCTGCAATTTC |
| MA75 | DIP5_S | TCAAAATTTGCTTATGTCGC |
| MA76 | DIP5_A | AGACAGGCAGGCCAATATAC |
| MA77 | TPS2_S | ATCAATGGGGCAACTACG |
| MA78 | TPS2_A | CGTACCAGCACTTTGGAAG |
| MA79 | GCD10_S | GACCTCAGGTTTTTAGCACC |
| MA80 | GCD10_A | TCCGATACAGGTTCAGGAG |
| MA81 | SPB4_S | ACTCGAGAGAAAGGAAAAGATG |
| MA82 | SPB4_A | GATAGCTTTGCTGGAAACTTTC |
| MA83 | YOL014W_S | TCTACTTGGCATGGTGTCC |
| MA84 | YOL014W_A | CATATTCGTTGGCTTCAGTG |
| MA85 | GFD2_S | ACCCTGCATTTGTTCATG |
| MA86 | GFD2_A | TCGATGGAGATGGCATAC |
| MA87 | DHR2_S | CTATAGGGATGCCAGACAGG |
| MA88 | DHR2_A | CATTTCTGGCATATCCCTTC |
| MA89 | GDH1_S | TTCTGGTTTAGAAATGGCAC |
| MA90 | GDH1_A | TTGACCAAAGATGGCAAG |
| MA91 | CLB6_S | GCATTGAAACAAGGAACATG |
| MA92 | CLB6_A | AAAAGTTTCATCCCATTTGG |
| MA93 | TOS6_S | ATGTTACCACCACCCCAC |
| MA94 | TOS6_A | CCGACGTATGTGCTGATC |
| MA95 | PRM7_S | CGTTCAATCAACTGCTTCC |
| MA96 | PRM7_A | CGTTAGTAGTGGTGGTCGTG |
| MA97 | ZRT1_S | GTGTTTTGGATGCCATTTC |
| MA98 | ZRT1_A | AAAGCCATGATACCAGCAC |
| MA99 | CRC1_S | TTGAACGTGTGTCTTGCTG |
| MA100 | CRC1_A | CCCTTGATACCACCTCTTTG |
| MA101 | CIT1_S | GTTGGTCTCCACCATTTATG |
| MA102 | CIT1_A | TGGCAACACCAAACAATAC |
| MA103 | RME1_S | GTCCCATAGAGCAATGTCC |
| MA104 | RME1_A | AAATGGGCAATTCAGTCC |
| MA105 | UGA1_S | GGGTTTGCAGAAGAAATACC |
| MA106 | UGA1_A | GACTGCACATCCACCAAC |
| MA107 | PCL9_S | AATACCAACCGTCCCTACC |
| MA108 | PCL9_A | ACTGGTGAACTTCCCATTG |
| MA109 | GPG1_S | TCGGAGAGTTGCACACAC |
| MA110 | GPG1_A | TTCATTGAAAGCACATCCC |
| MA111 | CHS2_S | AATTGTGATGATTTGGATGC |
| MA112 | CHS2_A | ACAAAAAGGCCATGGAAC |
| MA113 | PCL1_S | TGGCACGAGTCCTATCAG |
| MA114 | PCL1_A | CTGTGTTGTTCGCTATGTTG |
| MA115 | CTS1_S | ATATGCGGCTGGTAAATTG |
| MA116 | CTS1_A | AGTCGCCGGAGTCATAAG |
| MA117 | EGT2_S | GCCGAGCACACAAATTTAG |
| MA118 | EGT2_A | TGGAACTTGTTGGCAATG |
| MA119 | CLN1_S | TGCCATAAGCGTAAGCAG |
| MA120 | CLN1_A | GTCATAAATTTGGCACGTTG |
| MA121 | CLN2_S | AGCAATAACGCAACCAATG |
| MA122 | CLN2_A | TTTATGGTCCCAGTTGGC |
| MA123 | DSE1_S | ACCGGTAAAATAATCGATGG |
| MA124 | DSE1_A | ATTTGCTGAAGAACAATCCC |
| MA125 | DSE2_S | CGTCATCGTCTTCCACTTC |
| MA126 | DSE2_A | AGACGGATATCGATTGCG |
| MA127 | DSE3_S | AGTTGCACAAACATCATCG |
| MA128 | DSE3_A | CAAAGCCCTATCCTCGTC |
| MA129 | DSE4_S | ATTGGTTCAACGGTCATTC |
| MA130 | DSE4_A | TCAAGTCACCCCTCAATTC |
| MA131 | SCW11_S | GAAACTGCCGGTACATTTG |
| MA132 | SCW11_A | TTTGGACTGCAGTAATTTGG |
| MA133 | HXT1_S | CTACGGTTACGTTTTCATGG |
| MA134 | HXT1_A | CTTGGATACTGGAACCCAG |
| MA135 | FBP1_S | CCTTTTCGCATACCCTTG |
| MA136 | FBP1_A | TCATGGATATGACTTGGCAC |
| MA137 | AQR1_S | GGTGCAGTTTGAGTACCATC |
| MA138 | AQR1_A | ATCCTCCCTCCACTTCATAC |
| MA139 | JEN1_S | CATCTTCACATTTGCTTGTG |
| MA140 | JEN1_A | CATTCCGTCTTTTGTTCAAC |
| MA141 | YIG1_S | CATGCTTCAGGAACAGGAC |
| MA142 | YIG1_A | CCGTATTTGGCCCTAATG |
| MA143 | MIG2_S | CACCACCAAGACTAGGAGG |
| MA144 | MIG2_A | TACGCCTGACAATTTCTTTC |
