## Supplementary material for "TORC1 regulates the transcriptional response to glucose and developmental cycle via the Tap42-Sit4-Rrd1/2 pathway in *Saccharomyces cerevisiae*": Table S6

**Table S6**. Lists of genes in the 10 categories reported in ‘Spore germination in *Saccharomyces cerevisiae*: global gene expression patterns and cell cycle landmarks’ (Genome Biol. 2007; 8(11): R241). Comparative expression analysis of these 10 categories in our RNA-seq data is presented in Figure S2.

| Protein synthesis | *RPL19A,RPS8A,RPL17A,RPS18B,RPS10A,URP1,CRY1,YS29B,RPL43A,YDL082W,YDL083C,*  *SOS2,SOS1,RPS18A,RPS13C,RPL45,RPL15A,RP51B,YDR450W,RPL27B,RPL35B,RPL15B,*  *RPS24EA,RPS8B,RPL17B,RPS26B,RPL32,RPL30A,RPL6A,CYH2,SUP44,SSM2,RPL9A,*  *RPS31A,YGR034W,RPL16A,RPS28A,RPL30B,YST1,RPL14B,URP2,RPL4A,RPL27,MAK18,*  *RPS7A,RPL5A,YIL052C,RPL13,UBI1,RPS25B,TIF2,YJL177W,RPS24A,RPS5,RPS7B,*  *RPL14A,YKL056C,RPS27A,RPL17,RPS25,TIF1,UBI2,RPL4B,RPL13A,YST2,YLR061W,*  *GRC5,UBI3,RPL35A,RPS33B,YLR325C,RPS31,YLR388W,RP10A,RPL16B,YML024W,*  *YML026C,RP10B,YL16A,BEL1,YMR142C,YMR242C,RPL9B,RP23,YNL096C,RPL41A,RPS3,*  *SSB2,RP28B,RPS16A,RPLA2,RPS21,RP28A,RPS16B,RPL25,TCM1,RPS30,RPS33A,*  *RPL37B,YOR293W,RPL18A1,RPS12,EGD1,YPL079W,YPL090C,RPL37A,SSM1,YPR102C,*  *RPS28B,YBR084CA,YER056CA,YFR031CA* |
| --- | --- |
| rRNA processing | *YDR101C,YDR496C,ROK1,YGR103W,YGR145W,YGR245C,DRS1,PWP1,YLR222C,DBP9,*  *YLR409C,YML093W,HAS1,YNL132W,YNL174W,YNL182C,YOR206W* |
| Gluconeogenesis | *YCR010C,ICL1,YFL030W,YGR067C,HXT5,YIL057C,MBR1,YKL187C,JEN1,PCK1,IDP2,*  *FBP1,CYB2,YMR107W,YMR206W,MLS1,YNL194C,YNL195C,GAC1,LEE1,PXA1,YPR030W* |
| Stress | *YDL204W,TPS2,YGL037C,STF2,CTT1,SOL4,GRE3,OM45,YJR096W,YKL091C,TFS1,*  *YLR251W,YLR252W,TSL1,YML128C,PGM2,YMR250W,YNL274C* |
| TCA sub-cycle | *CIT2,ACO1,IDH1,CIT1,YOR135C,IDH2,YPL087W,YPL135W,PEP4,YPR002W* |
| Oxidative phosphorylation | *PET9,COR1,ATP1,ATP3,YBR183W,YBR230C,ATP16,COX9,INH1,SDH4,ATP5,ATP17,*  *QCR7,RIP1,YER053C,COX15,QCR6,COX4,COX13,CBP4,YGR182C,QCR9,COX6,QCR8,*  *CYC1,MIR1,ATP2,ATP7,MDH1,HAP4,SDH3,SDH1,MCR1,SDH2,COX12,YLR294C,*  *ATP14,COX8,NDI1,COX7,PBI2,COX5A,POR1,YNL100W,CIT1,CYT1,ATP4,ATP15,*  *YPR020W,QCR2,YHR001WA* |
| Proteasome subunits | *PRE7,YBR062C,YBR173C,YTA5,RPN5,RPN4,YTA2,RPN8,PRE1,SUN2,PUP3,MPR1,*  *YFR010W,PRE4,NIN1,SCL1,SUG1,UFD1,PRE9,PHB2,PUP2,ARC15,PRE3,CAP1,*  *SBA1,YTA3,YKT6,YKR011C,YLR387C,YLR421C,GLO1,PRE8,PRE5,YNL155W,PRE6,*  *CRL13,RPN7,PRE10,PRE2,RPN6* |
| Mating | *PRM9,YAR033W,FUS3,YBL062W,FIG1,YBR156C,YBR158W,YBR223C,YBR225W,*  *YBR226C,FUS1,KAR4,YCL074W,YCL075W,YCL076W,RVS161,FIG2,PCL2,RDI1,*  *AFR1,YDR124W,ECM18,YDR241W,YDR249C,PAM1,YDR309C,YDR340W,STE14,MFA1,*  *YER187W,STE2,YFL027C,YFL047W,AGA2,YGL052W,PRM8,IME4,YGL223C,GPA1,*  *STE12,YHR097C,CHS7,PRM2,YIL060W,YIL080W,YIL082W,YIL083C,PRM5,*  *YJL107C,PRM10,FAR1,ASG7,PGU1,GFA1,PGM1,PMU1,HYM1,YKL221W,KTR2,*  *YLR042C,MID2,SST2,PRP39,PRM6,KAR5,CIK1,FUS2,YNL042W,MSG5,INP52,*  *CHS1,YNL208W,PRM1,ERG24,AGA1,YOL095C,YOR129C,YOR343C,PRM4,PRM3,*  *YPL193W,KAR3,YML048WA,YIL082WA,YMR304CA* |
| Cell-cycle (G1) | *RFA1,SEN34,HTB2,HTA2,YBL009W,POL12,HHF1,HHT1,YBR070C,YBR071W,RDH54,*  *RFC5,POL30,YBR089W,YCL022C,YCL024W,YCL061C,HCM1,MCD1,YDL018C,DUN1,*  *YDL163W,CDC9,ASF2,MSH6,PDS1,HTB1,HTA1,YDR279W,GIN4,YDR528W,MNN1,*  *PMI40,RNR1,RAD51,SSU81,ADK2,SMC1,CLB6,YGR151C,RSR1,YGR221C,YHR110W,*  *YHR127W,SPO16,YHR154W,YHR173C,IRR1,YIL132C,SRO4,SMC3,HPR5,ASF1,RFA3,*  *YJL181W,SWE1,POL32,PRI2,MIF2,HSL1,YKL108W,RAD27,YKR077W,YKR090W,KIM2,*  *SPA2,YLL022C,STU2,YLR049C,CDC45,YLR183C,TUB4,SPH1,YOX1,OGG1,CTF18,*  *YMR144W,SPT21,CLN1,HHF2,HHT2,PMS1,POL1,SPC98,YNL166C,BNI4,POL2,YIF1,*  *TOF1,RFA2,YNR009W,YOL007C,YOL017W,MSH2,BUB3,DHS1,CDC21,YOR114W,*  *YOR144C,NIP29,HHO1,RAD53,SVS1,IPL1,BBP1,CLN2,YPL267W,RLF2,CLB5,DPB2* |
| Cell cycle G2/M | *KIN3,YBL032W,CHS2,PHO3,CDC47,BUD3,YDR033W,SWI5,PMA1,ALK1,CDC20,DBF2,*  *CLB1,WSC4,MOB1,YIL158W,YJL051W,BUD4,YKL130C,YLR084C,ACE2,YLR190W,*  *YML034W,SUR7,YML058W,YML119W,CDC5,YMR032W,YMR215W,YNL057W,YNL058C,*  *YOL070C,HST3,YOR315W,YPL141C,KIP2,IQG1,CLB2,YCR024CA* |
